# Local mechanical stimuli shape tissue growth in vertebrate joint morphogenesis

**DOI:** 10.1101/2021.08.28.458034

**Authors:** Ester Comellas, Johanna E Farkas, Giona Kleinberg, Katlyn Lloyd, Thomas Mueller, Timothy J Duerr, Jose J Muñoz, James R Monaghan, Sandra J Shefelbine

## Abstract

The correct formation of synovial joints is essential for proper motion throughout life. Movement-induced forces are critical to creating correctly shaped joints, but it is unclear how cells sense and respond to these mechanical cues. To determine how mechanical stimuli drive joint morphogenesis, we combined experiments on regenerating axolotl (*Ambystoma mexicanum*) forelimbs with a poroelastic model of bone rudiment growth. Animals either regrew forelimbs normally (control) or were injected with a TRPV4 agonist to impair chondrocyte mechanosensitivity during joint morphogenesis. We quantified growth and shape in regrown humeri from whole mount light sheet fluorescence images of the regenerated limbs. Results revealed statistically significant differences in morphology and cell proliferation between groups, indicating that mechanical stimuli play a role in the shaping of joints. We simulated local tissue growth in a finite element model with a biological contribution to growth proportional to chondrocyte density, and a mechanical contribution to growth proportional to fluid pressure. Computational predictions agreed with experimental outcomes, suggesting that interstitial pressure driven from cyclic mechanical stimuli promotes local tissue growth. Predictive computational models informed by experimental findings allow us to explore potential physical mechanisms involved in tissue growth to advance our understanding of the mechanobiology of joint morphogenesis.

## 1 Background

The shape of a synovial joint is critical to its functionality in movement and locomotion. Joint morphogenesis in the developing vertebrate limb bud follows a well-known sequence of events [1]. First, the mesenchymal cells forming the early limb bud differentiate into chondrocytes, except for those in the interzone, where the future joint will appear. Through a process known as cavitation, the skeletal rudiments are physically separated and the synovial cavity is formed. After cavitation, chondrocyte proliferation and matrix production in the rudiment result in growth and final joint shape. Movement-induced mechanical stimuli condition the correct formation of joints throughout this morphogenetic stage [2,3]. Yet, how motion and biophysical forces influence joint shape is not fully understood to date [4,5]. Insights into how chondrocytes proliferate and regulate joint shape in response to mechanical stimuli during morphogenesis has application in the study and treatment of joint deformities [5].

Animal studies using immobilised chicks [6–10], reduced-muscle and absent-muscle mice [11–13], and paralysed zebrafish larvae [14] have shown that reduced and restricted muscle contractions during embryonic development results in skeletal abnormalities, including alterations in joint shape. Elucidating the role of motion in joint development is challenging in animal models that develop in ovo or in utero [3]. An animal model that allows rigorous control of the biophysical environment during joint morphogenesis will further our understanding of how mechanical stimuli are linked to cell proliferation and tissue growth. Axolotl salamanders (*Ambystoma mexicanum*) regenerate limbs throughout life by recapitulating developmental processes. Regenerating axolotl limbs undergo stereotypical patterns of gene expression and cell differentiation that resemble mammalian joint development [15,16]. Their limbs are morphologically similar to human limbs, with elbow joints comparable in cellular composition and skeletal structure to mammalian synovial joints [17,18].

Joint morphogenesis in vertebrates is driven by the proliferation and subsequent hypertrophy of chondrocytes that form the bone rudiments. Chondrocytes respond to mechanical stimuli such as changes in osmotic pressure, cellular stretch, or fluid shear [19]. Ion channels, integrin signalling, and the primary cilia are all known mechanosensors that initiate intracellular signalling cascades ultimately resulting in the transcription, translation, and/or molecular synthesis that leads to cartilage tissue growth [19–21]. In vitro studies have shown that the transient receptor potential vanilloid 4 (TRPV4) channel is possibly a key transducer of biophysical stimuli to regulate cartilage extracellular matrix production [22–24]. TRPV4 activation in chondrocytes has been linked to osmolarity changes in in vitro studies [25,26]. Recent studies have shown it also responds to physiologic levels of strain loading [27,28], although there is also evidence to the contrary [29,30].

To identify the specific mechanical stimuli influencing joint shape, computational models can help decipher the role of biophysical stimuli in tissue growth and joint morphogenesis. Techniques like finite element analysis (FEA) are specially suited to studying the mechanics of morphogenesis. They allow for the quantitative, unbiased testing of the biophysical mechanisms that might be regulating and controlling morphogenesis [31,32]. A few studies have used FEA to examine how changes in mechanical loading affect joint morphogenesis [33–36]. These models demonstrated shape changes based on generic joint shapes and idealised loading conditions in two dimensions. The computational models assume that dynamic hydrostatic compression promotes cartilage growth, which is in line with experimental studies that have shown an increase in extracellular matrix production with cyclic compression [37–41]. Yet, these numerical studies use a static approximation via linear elasticity. As such, they are unable to intrinsically capture the effects of dynamic loading on cartilage, including the fluid flow and extracellular pressure to which cells likely respond. To better comprehend how local mechanical stimuli drive the shaping of the joint, we must model the tissue as a poroelastic medium, which incorporates a fluid component to account for the dynamic changes in pressure and velocity of extracellular fluid present in cartilage.

The goal of this study was to determine the effect of limb motion on joint morphology, and identify potential mechanisms by which mechanical loading is translated into chondrocyte proliferation and unequal tissue growth that results in joint shape. Experiments on regenerating axolotl limbs provided information on how altering chondrocyte mechanosensitivity affects the shaping of the joint. Predictive computational models informed by the experimental findings allowed us to explore potential physical mechanisms influencing joint morphogenesis.

## 2 Effect of impaired mechanosensitivity on regrowing axolotl elbow joints

To determine the effect of local mechanical stimuli on tissue growth during joint morphogenesis we restricted the ability of cells to respond to mechanical stimuli in regrowing axolotl forelimbs using the TRPV4 agonist GSK1016790A. Most known genetic disfunctions of the TRPV4 channel resulting in skeletal dysplasias are related to a gain of function [42,43]. The lack of regulation of intracellular calcium ions induced by the chemical activation of TRPV4 channels means that the chondrocytes lose their mechanosensitivity and are effectively unable to detect and respond to mechanical stimuli [44,45].

### 2.1 Axolotl experiments

Larval animals (3-5 cm) were bilaterally amputated just proximal to the elbow joint. GSK1016790A was reconstituted in dimethyl sulfoxide (DMSO) and injected intraperitoneally at 50 μg/kg at 22 days post amputation (dpa, n=6). Control animals (n=6) were injected with 50 μg/kg DMSO. Injections were repeated at 48-hour intervals. At 32 dpa, all animals were injected intraperitoneally with 5-Ethynyl-2’-deoxyuridine (EdU) and L-Azidohomoalanine (AHA). Limbs were collected 18 hours later, fixed and stained.

We imaged nascent macromolecule synthesis in the regenerated forelimbs with light sheet fluorescence microscopy following the whole-mount click-it-based technique in Duerr et al. [46]. EdU is incorporated into newly-synthesised DNA, which allowed for the quantification of cell proliferation through EdU-positive nuclei segmentation. AHA enabled visualizing chondrocyte protein translation, most likely extracellular matrix, which provided a well-defined outline of the bone rudiment’s perichondria. Quantification of 3D shape was then possible through the analysis of the humerus outline (Fig. 1A).

**Figure 1:**
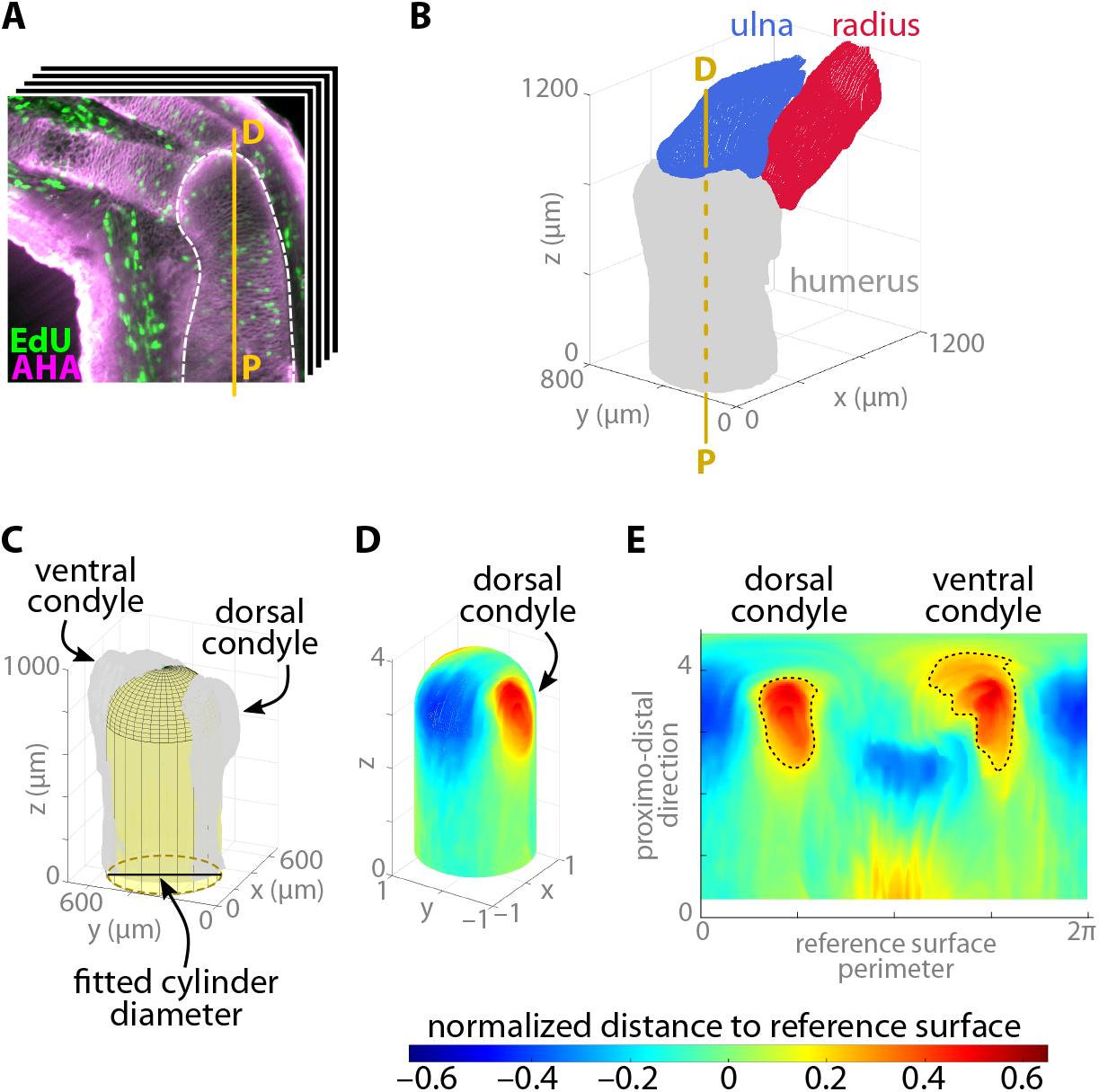
Overview of the experimental data analysis pipeline. (A) 3D image stacks of the axolotl elbow were acquired using light sheet fluorescence microscopy following the technique in Duerr et al. [46]. Stacks were cropped and rotated to vertically align the proximo-distal (P-D) axis of the humerus. EdU staining (green) was used to identify the proliferating cell nuclei within the humerus bone rudiment. AHA staining (magenta) allowed for segmentation of the bone rudiments. (B) The segmented bone rudiment surfaces were oriented in 3D space based on the longitudinal axes of the humerus and ulna. An elbow of an exemplary left forelimb is shown. (C) A reference surface (yellow) consisting in a cylinder with a hemispherical cap was fitted to the aligned humerus surface (grey). (D) The perpendicular distance from the reference surface to the humerus surface was mapped onto the reference surface. Both the mapped distance and the reference surface dimensions were normalized with the fitted cylinder diameter. (E) The mapped values were flattened out (unwrapping the reference surface) to obtain a 2D standardised humerus surface representation, where red indicates a protuberance (the condyles) and blue shows the concavities of the original 3D surface. The normalized areas and normalized volumes of the dorsal and ventral condyles were then extracted for analysis. A threshold value of 0.2 was considered to define the contour of the condyles (dashed line).

SI Appendix, Fig. S1A illustrates the timeline of the experiments and SI Appendix, Fig. S1B shows an example of the animal size used. Injections started at 22 dpa, which is roughly when joint cavitation occurs in regenerating limbs in 3-5-cm-sized animals, and continued throughout the joint morphogenesis stage of the joint formation process until 30 dpa. SI Appendix, Fig. S1C shows a central slice of a 3D image stack obtained for an exemplary control elbow. All light sheet images were acquired using a Zeiss light sheet Z.1 microscope paired with Zen software.

### 2.2 Experimental data analysis

We segmented the bone rudiments, identified the proximo-distal longitudinal axis of the humerus and ulna through computation of the minimum principal axis using the Fiji plugin BoneJ [47], and then aligned all limbs in 3D space (Fig. 1B). The alignment process included mirroring of right limbs so that all limbs had the dorsal and ventral condyles in the same relative position in space.

A cylinder was fitted to the aligned humerus surface using the Matlab [48] File Exchange function ‘cylinderfit’ (a regression modelling tool), and a hemispherical cap was placed on top to create the reference surface (Fig. 1C). These were shifted vertically upwards until the hemispherical cap was tangent to the distal end of the humerus surface. The distance from the reference surface to the humerus surface was mapped onto the reference surface and normalized using the fitted cylinder diameter (Fig. 1D) to account for animals of different size. We quantified dorsal and ventral condyle shapes and sizes based on the corresponding normalized areas and normalized volumes, respectively, which were extracted from the 2D standardised representation of the humerus surface (Fig. 1E).

To quantify proliferating cells, we manually generated a small training set to train the deep learning algorithm Stardist3D [49], which was used to identify the EdU-stained cell nuclei in the 3D image stack. Memory limitations in Stardist3D required splitting the original image stack into smaller substacks for processing. The Fiji plugin 3D Objects Counter [50] was used on the cell nuclei masks produced by Stardist3D to identify proliferating cell positions and volumes. The data was then regrouped and the whole set was masked with the humerus bone outline segmented from the AHA channel. The alignment data obtained in the humerus shape analysis process was used to align the cell nuclei in 3D space. Outliers were removed based on cell volume and we used a fixed-length cut-off to ensure quantification of cell proliferation was performed in an equivalent humerus volume across different limbs.

SI Appendix, Fig. S2 provides a visual summary of the complete workflow, which was implemented using a combination of Fiji [51], the ZeroCostDL4Mic implementation of Stardist3D [52] and a customised code in Matlab [48].

### 2.3 Experimental results

Our results reveal significant differences in cell proliferation and shape between the humeri of the control group and the mechanosensitively-impaired (GSK101) group. The mean value of the proliferating cell count in the humeri of the control group is fourfold that of the GSK101 group (p-value*<*0.001, Fig. 2A) A central slice of each EdU-stained humeri (Fig. 2D) and the 3D distribution of the cell nuclei identified in two representative humeri of each group (Fig. 2E) depict the differences in cell proliferation between groups. However, the diameter of the humeri shaft is similar for both groups (Fig. 2B).

**Figure 2:**
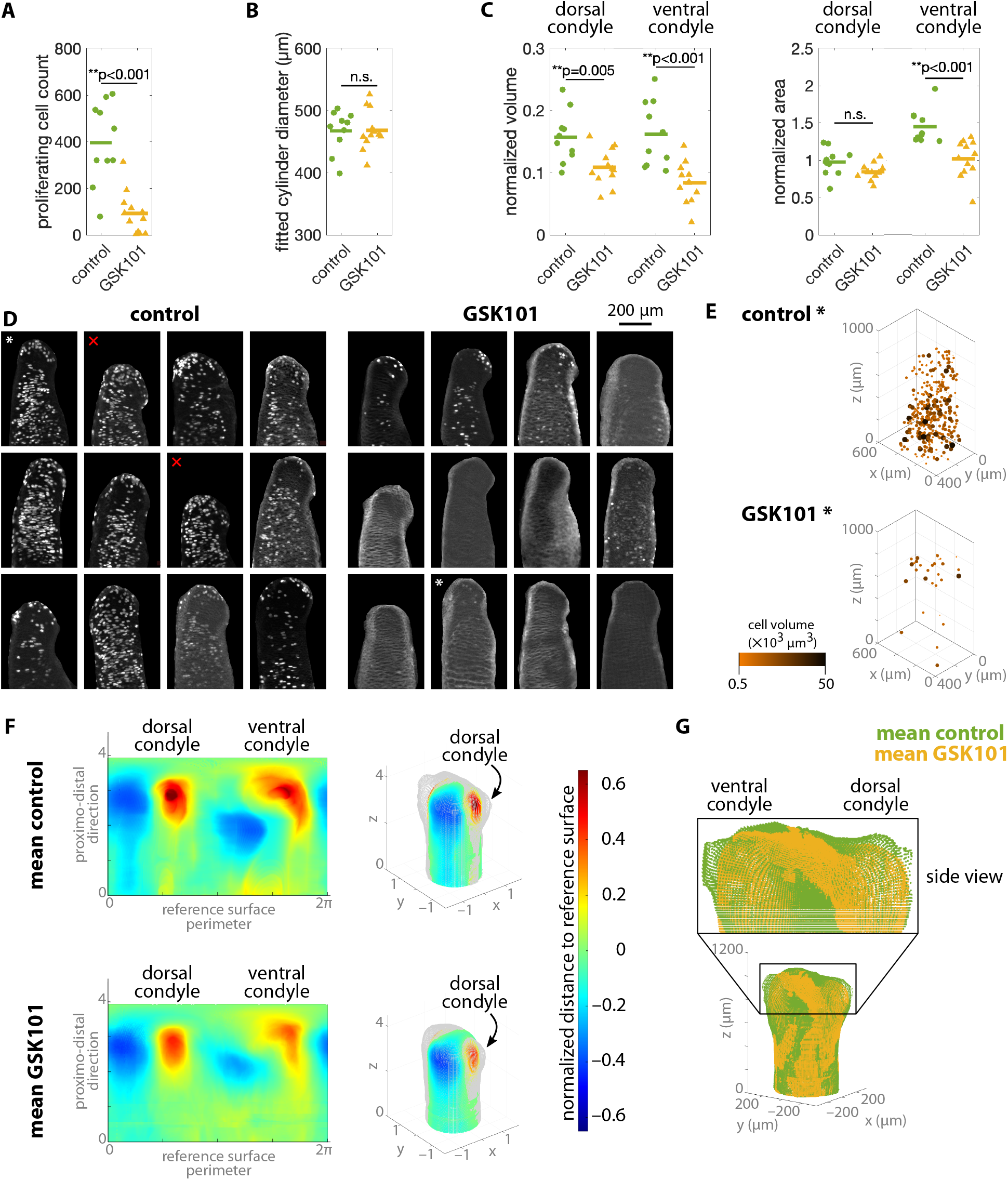
Quantification of humerus growth and shape in regenerated axolotl humeri. (A-C) Results of the statistical analysis on the data points. Measurements were obtained following the methodology outlined in Fig. 1. Control group (n=10) and mechanosensitively-impaired (GSK101) group (n=12); For group comparison in A: Kruskal-Wallis test; For B and C: one-way ANOVA. Shapiro-Wilk test was used to test for normality. (D) EdU-stained masked humeri. The maximum intensity projection of 20 central slices in each humerus is shown. All images have the same 200 μm scale bar (top right). The two control humeri marked with a red cross were excluded from the 3D shape analysis (A-C) because they were too short to be aligned with the methodology developed in this study. (E) 3D cell nuclei positions for a representative humerus of each group (marked with an asterisk in (D)). (F) Mean 2D surface maps for the control (top row) and GSK101 groups (bottom row). The corresponding mean normalized humerus surface (in grey) is recovered for both groups in 3D space. (G) The mean humerus surface for the control and GSK101 groups are aligned for comparison. A diameter of 450 μm is used for both.

Regarding humeri shape, the normalized volumes of both dorsal and ventral condyles are larger for the control group than the GSK101 group (p-value=0.005 and *<*0.001, respectively). The normalized areas of the ventral condyles in the control group are also larger (p-value*<*0.001), while no significant difference was found for the normalized areas of the dorsal condyles (Fig. 2C). The mean humerus surfaces for the control and GSK101 groups allow for the reconstruction of the corresponding mean humerus 3D surfaces (Fig. 2F). These are overlaid for comparison (Fig. 2G), showing more prominent condyles in the mean control humerus. Individual 2D surface maps used to compute the means are provided in SI Appendix Fig. S3. To compute the mean 2D surface maps (Fig. 2F, first column), we previously aligned the individual 2D surface maps (SI Appendix, Fig. S3) based on the position of the dorsal condyle centroids.

## 3 Computational predictions of joint morphogenesis

We created a finite element model of a regenerating humerus to explore potential movementinduced mechanical stimuli as drivers of tissue growth. Bone rudiment is composed of cartilage tissue, which has a water content of roughly 80% by volume of tissue mass [53]. The mechanism for transduction of mechanical forces in tissues is not completely understood, but fluid flow is known to play an important role [19]. Poroelastic theory is commonly used in finite element models of cartilage response to loading [54–57] because it can explicitly capture the fluid flow effects.

### 3.1 Modelling cartilage tissue growth within a poroelastic framework

The biphasic approach defines tissue as a mixture of an elastic solid skeleton with free-flowing fluid circulating within its pores. In cartilage, the fluid can be assimilated to the interstitial fluid in the tissue, i.e. water and dissolved ions, growth factors and other molecular components. The solid component represents the proteoglycans and collagen of the extracellular matrix (ECM) and chondrocytes. Chondrocyte proliferation and ECM production in cartilage can then be modelled through continuum growth of this solid phase.

The deformation of the solid component is characterized by its displacements ***u***_*S*_ while the fluid behaviour is defined by the pore pressure *p*. The governing equations required to solve the problem for the two unknowns, ***u***_*S*_ and *p*, are the linear momentum and mass balance equations. These introduce the constitutive equations of the solid and fluid components, respectively. The solid behaviour is characterized by the Kirchhoff stress tensor ***τ*** while the fluid flow is defined by the volume-weighted seepage velocity ***w***, which is the relative velocity of the fluid with respect to the deforming solid. For simplicity, we considered a neo-Hookean hyperelastic model for the solid part and a Darcy-like law for the fluid.

Tissue growth is modelled via the multiplicative decomposition of the deformation gradient tensor ***F*** that characterizes the solid component deformations, which include both the deformation and the growth due to loading (SI Appendix, Fig. S4). For simplicity, the growth tensor is assumed to be volumetric and proportional to the growth stretch *ϑ*. Following a common approach in the field [32–34,58] we consider growth rate to be a sum of biological and mechanical contributions, denoted respectively by 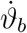 and 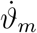.

The biological contribution represents the intrinsic morphogenetic biological factors that globally mediate tissue growth. Similar to past studies of joint morphogenesis [33,34], we assumed it is proportional to chondrocyte density in the bone rudiments. However, unlike these studies, our experimental measurements of chondrocyte density in a regenerating axolotl humerus revealed an approximately constant value throughout the bone rudiment (SI Appendix, Fig. S5). Therefore, we defined a constant biological growth stretch rate 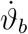 in time and space, within the humerus geometry and throughout the whole simulation time period.

The mechanical contribution is a function of the selected mechanical stimulus locally driving tissue growth. Mechanical loading is known to modulate the synthesis of ECM in chondrocytes. Collagen and aggrecan production, the main components of ECM in cartilage, depends on the magnitude, duration and type of loading. In particular, in vitro experiments have shown that cyclic compression promotes ECM production while static loading either has no effect on collagen and aggrecan levels, or inhibits cartilage growth [37–41]. Based on this experimental evidence, past models considered (compressive) hydrostatic stress as a driver of mechanical growth [33–35]. Even with the simplifying assumptions considered and generic joint shapes used, Giorgi et al. [34] could predict anatomically recognisable joint shapes based on different starting joint configuration and applied movements. Their results indicate that hydrostatic stress could be mediating tissue growth in response to mechanical load. However, models to date used a single-phase elastic material to represent tissue behaviour and, hence, were unable to inherently distinguish between dynamic and static loading effects. Our poroelastic model overcomes this limitation and, for the first time, we are able to define mechanical growth proportional to a dynamic variable linked to the movement-induced fluid flow. We selected pore pressure of the fluid component, a hydrostatic measure akin to the hydrostatic stress used in past models, as the mechanical stimulus.

The discretized governing equations and continuum growth model were implemented in the open source finite element library deal.II [59]. The code used in this study is an extension of the poro-viscoelastic numerical framework in [60]. Growth was implemented following the algorithm in SI Appendix, Fig. S6. Quadratic shape functions were used to approximate the solid displacements, linear shape functions were used to approximate the pore pressure, and a quadrature of order 3 was considered in all the simulations. Further details of the poroelastic formulation, the growth model and their numerical implementation are provided in SI Appendix, Section.

### 3.2 A finite element model of joint morphogenesis

We generated a finite element (FE) model of a generic humerus bone rudiment after cavitation, i.e. at the start of the experiments, with the goal of predicting the grown humerus shape at the end of the joint morphogenesis stage. Given that our model is a tool to probe potential mechanisms of load mechanotransduction in joint morphogenesis, we strove to keep its parameters as generic as possible.

The geometry and loading conditions (Fig. 3A) were informed by experimental data from a regenerating axolotl forelimb just after joint cavitation. We segmented the bone rudiment shapes of a normally-regenerating forelimb at 17 dpa in a 3-cm-sized animal (SI Appendix, Fig. S7A).

**Figure 3:**
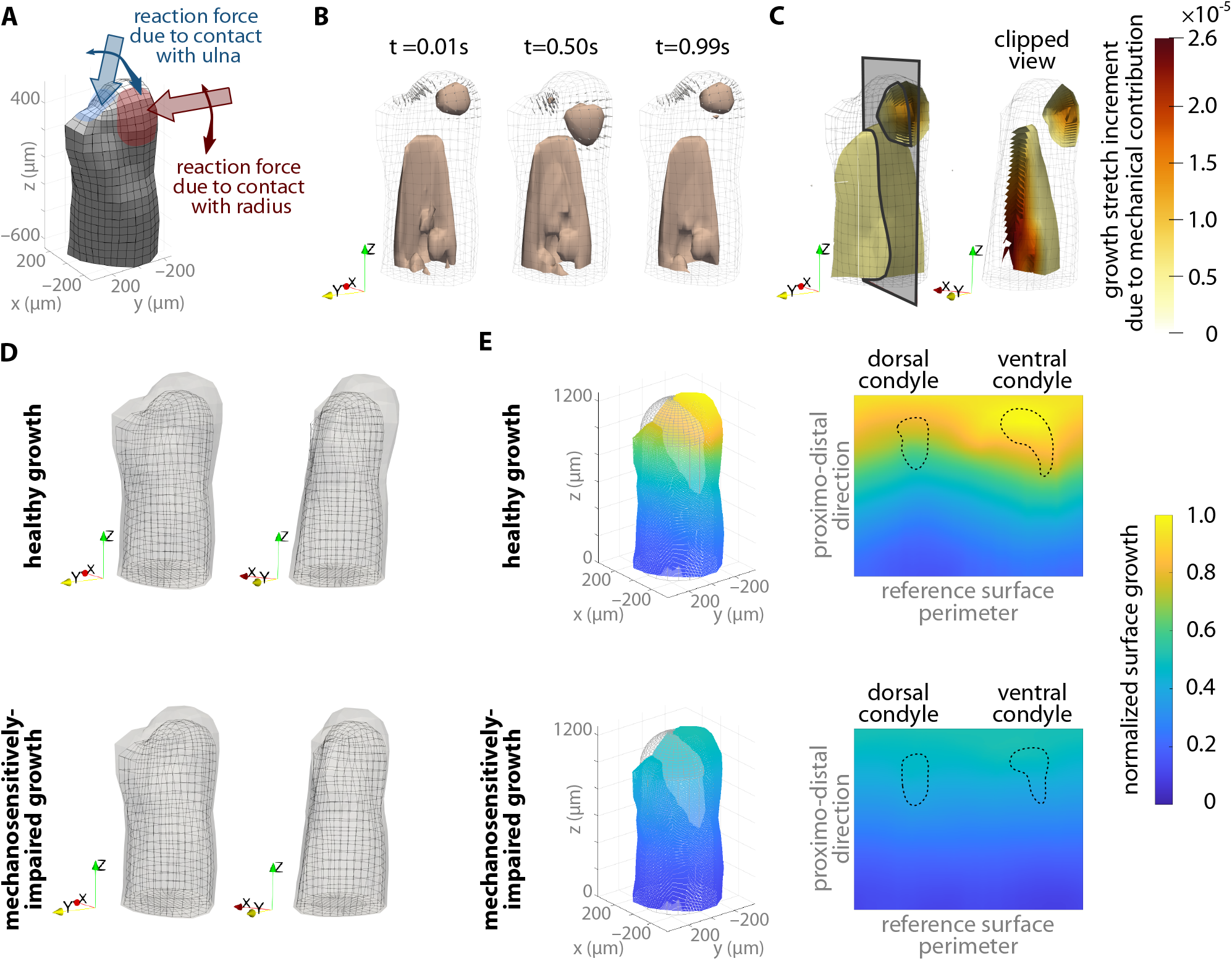
Computational predictions of joint morphogenesis considering pressure-driven local tissue growth. (A) Finite element model of the humerus. Loading to simulate a 1-second flexion-extension cycle of the elbow was applied as a sweeping motion together with a twofold increase in pressure load intensity at peak flexion. (B) Predicted pressure distribution in the whole humerus at the start, middle, and end of a flexion-extension cycle. Small grey arrows on the surface nodes represent the applied loading at each time point. A pressure contour of 1 kPa is shown for all. (C) Local tissue growth due to the mechanical contribution at the end of one cycle. Local tissue growth due to the biological contribution was constant and not shown here. (D) Grown humerus shape representing a healthy case (top) and a mechanosensitively-impaired case (bottom). Local tissue growth in the healthy case, comparable to the experimental control group, included both mechanical and biological contributions. The grown humerus shape for the mechanosensitively-impaired case, comparable to the experimental GSK101 group, had local tissue growth in response to the biological contribution only. Growth is scaled by a factor of 3600, representing a 1-hour period, and frontal (left) and side (right) views are shown for both. (E) Quantification of grown humerus shapes based on the normalized surface growth, mapped onto a reference surface like in the experimental data analysis, and then flattened into a 2D map of normalized surface growth.

A mesh was generated based on the smoothed-out surface of the segmented humerus with a total of 512 hexahedral elements. We scaled the geometry size to achieve a cross-sectional humerus size closer to the values identified in our experiments. Meshing of the geometry inevitably entails a slight loss of surface detail. We computed and visually compared the 2D surface maps of both the segmented geometry and the meshed geometry (SI Appendix, Fig. S7B) following a procedure analogous to the one used in the humerus 3D shape analysis. Comparison of 2D surface maps confirmed that the meshed surface retained the main characteristics of the original humerus.

Free-flow boundary conditions across all external surfaces except the bottom (proximal) one were set in the FE model. Vertical displacements of the bottom surface were fixed, and lateral displacements of nodes in the bottom surface were fixed (SI Appendix, Fig. S7C). These boundary conditions allowed for outward growth of the humerus shaft while avoiding spurious translations as well as the rotation of the whole bone rudiment.

The loading conditions applied (Fig. 3A), modelled a 1-second flexion-extension cycle of the elbow. The growth resulting from a single cycle was extrapolated for multiple cycles. Loading was applied as a pressure over a roughly circular surface representing the contact areas between the radius/ulna and the humerus. A sine-like loading profile over this area was considered, with the loading area sweeping over the humerus surface. The sweep path was estimated based on anatomical observations of the axolotl elbow joint. The value of the load profile changed throughout the cycle to mimic the effect of muscle contractions, reaching the maximum value for the peak flexion position. Load step increments of 0.01s were applied. We studied the effect of varying loading and boundary conditions on our computational results (SI Appendix, Fig. S8). In this way, we ensured the robustness of our computational setup to produce results from which to extract meaningful insights.

The material properties were either estimated from literature or based on an educated guess, except for the initial intrinsic permeability of the biphasic material. Preliminary simulations identified this parameter as having a considerable impact on the predicted patterns. Hence, we adjusted its value based on experimental stress-relaxation data obtained through nanoindentation tests on an axolotl forelimb (SI Appendix, Fig. S9). SI Appendix, Section provides further details of all model parameters.

### 3.3 Computational results

A regenerating humerus model based on local changes in fluid pressure (Fig. 3B) induced by an elbow flexion-extension loading cycle predicted a final humerus morphology that resembled our experimental observations of the control group (Fig. 3D, top row). When the mechanically-driven growth component (Fig. 3C) was removed, shape prediction resulted from constant volumetric biological growth only and was in accordance with the experimental observations of the mechanosensitively-impaired GSK101 group (Fig. 3D, bottom row). When the mechanically-driven growth component was included, local mechanical growth occurred in regions of high compressive pressure, which were observed underneath the surface load representing the radius contact area throughout the cycle. However, we did not observe an analogous pressure below the load representing the ulna contact area. Pressure was most pronounced at the posterior proximal part of the humerus shaft. Complete predicted patterns for pressure as well as other mechanical stimuli that were initially considered as potential drivers of the mechanical growth model are provided in SI Appendix, Figs. S10 and S11.

To quantify the differences between the healthy and mechanosensitively-impaired cases, we computed at each surface node the magnitude of the distance between the original surface and the grown surface, and normalized this measure with the maximum value of the two cases. We then mapped the resulting patterns onto a reference surface, fitted to the original surface mesh, and flattened it to obtain a 2D representation of normalized growth (Fig. 3E). The mapping procedure was analogous to the one used to obtain the 2D surface maps of the experimental humeri (Fig. 1C-E). In both predictions, humerus surface growth increased towards the distal portion of the bone rudiment, but the the healthy growth case resulted in larger values as well as a notably asymmetrical pattern. A larger surface growth was predicted in the area corresponding to the future ventral condyle for the healthy growth case (Fig. 3E, top row). The contour of the condyles from the corresponding mean experimental surfaces in Fig. 2F is shown on the 2D maps.

## 4 Discussion

### 4.1 Impaired mechanosensitivity during joint morphogenesis altered final humerus shape

Our analysis of the regenerating axolotl limbs revealed an altered humerus morphology for the mechanosensitively-impaired (GSK101) group (Fig. 2A-C). The mean 2D surface maps computed for each group (Fig. 2F) illustrate the main findings: the condyles of the control animals have larger normalized volumes than the mechanosensitivity-impaired group (darker shade of red in contour map). The shape of the ventral condyle, as measured based on the normalized area, is more affected by lack of mechanics than the dorsal condyle. We also analysed the shapes and sizes of the anterior and posterior concavities (blue regions in the 2D surface maps, SI Appendix Fig. S3) following an analogous procedure to the condyle measurements and did not observe significant differences between groups for any measurement. Taken together, this data seems to indicate that, when unable to sense and respond to mechanical cues during joint morphogenesis, the final humerus shape is affected.

The fact that the dorsal condyle and concavity areas are equivalent in both groups could also signify that these shape characteristics of the humerus were already present at the onset of the experiment; starting treatment with GSK101 sooner after amputation may result in more severe changes. We analysed a regenerating limb after cavitation (start of our experiments) for the purposes of developing the initial computational model. The 2D surface map obtained (SI Appendix, Fig. S7B, left) supports the notion that the basic humerus shape could be present already at this stage. The concavities seem to already be present and the dorsal condyle is clearly defined, similar in shape to those of the fully regenerated limbs in the experiments (SI Appendix, Fig. S3). However, the ventral condyle is barely discernible after cavitation. This implies that the concavities and dorsal condyle may form in the earlier stages of the joint formation process, which is likely why we found little change in their shapes.

Numerous studies have shown that chondrocytes have several separate but overlapping mechanotransduction pathways [27,29]. Other channels of the TRP family have been suggested to have load-associated effects in cartilage [61], but TRPV4 is undoubtedly the major regulator of mechanical and osmotic signal transduction in this family. The Piezo1 and Piezo2 channels have also been identified as key stretch-induced mechanotransducers in chondrocytes [62]. It would be interesting to see whether altering these other channels has effects on morphology similar to those seen in this study, to further explore the interrelated roles of each channel in cartilage mechanotransduction.

Alternative ways of blocking mechanics in developing joints have been used in the past to study the effect of mechanical stimuli on joint morphogenesis, namely muscle paralysis in chicks [6–10] and genetically-modified altered-muscle mice [11–13]. These studies also revealed morphological differences. Here, we used a TRPV4 agonist, which represents the clinical genetic deficits associated with abnormal skeletal development [63,64]. In addition, our 3D analysis of the humerus surface allows the assessment of shape changes that are not evident in more simple measures used in the past, such as cross-sectional outlines or linear anatomic measurements like humeral head width.

### 4.2 More prominent condyles and increased chondrocyte proliferation were not associated with larger humeri

The substantial reduction in cell proliferation of the GSK101 group (Fig. 2A, D and E) did not result in smaller humeri sizes (Figure 2B). To ensure that the similar humeri fitted diameter values between the two groups were not due to an insufficiently sensitive measurement method, we computed additional metrics using an alternative methodology (SI Appendix, Section). All measures of humeri shaft size computed indicted there were no significant differences between the two groups.

This apparent discrepancy between amount of cell proliferation and humeri size could be due to the axolotl long cell cycles, which have been recorded to be up to 88 hours in regenerating tissues [65,66]. Then, throughout the 10 day experimental treatment, few complete cycles would have occurred. Considering that proliferating cells were only a relatively small percentage of total chondrocytes in the bone rudiment, and the small amount of cell cycles completed, maybe the total amount of cell proliferation was not enough to produce actual changes in bone rudiment size. In addition, our quantification of cell proliferation corresponded to an 18-hour window at the end of the experiment, which may not be representative of the complete treatment period of 10 days.

Yet, we identified a decrease in condyle normalized volumes and in the ventral condyle normalized area for the GSK101 group, which raises the question whether reduced cell proliferation could be related to altered humerus shape. condyle cartilage growth has been linked to cell proliferation in developing chick knee joints [8]. Interestingly, regulation of chick embryo limb growth in response to motility was linked to cell proliferation only in specific growth plates in a separate study [9]. In the present study, we did not observe proliferation localized to the distal part, on the contrary it was seemingly homogeneously distributed (Fig. 2D and E). Therefore, a direct link between cell proliferation location and localized tissue growth could not be made based on our experimental observations. It is highly likely that the specific proteins that contribute to tissue growth, such as extracellular matrix proteins, could lead to increased localized growth. Alternatively, directional cell growth, independent of cell proliferation, could lead to differential growth in particular areas of the tissue.

Injections in experiments started at 22 days post amputation (dpa) as this was the estimated time of joint cavitation in the regenerating forelimbs for animals 3-5 cm in size. We analysed a 17 dpa regenerated forelimb of a 3-cm-sized animal to develop our finite element model (SI Appendix, Fig. S7), which revealed that the bone rudiments were fully formed and separated, and the humerus already had a rudimentary shape, including a defined dorsal condyle. It is possible that our experiments in fact targeted the final stages of joint morphogenesis, and the bone rudiment was already close to its final size from the start of the injections. Given that regenerating limbs grow outwards from a limb bud that is created from a fully-grown stump, the proximal portion of the humerus is already correctly sized, while the joint is undergoing morphogenesis in the distal part. In contrast, during development, one would expect the joint to form as bone rudiments around it are also growing in size. Considering all the above, there are multiple possibilities as to why no difference was observed in humerus shaft size between the two experimental groups.

### 4.3 Local fluid pressure may promote tissue growth during joint morphogenesis

Our experimental results provide additional confirmation that mechanical forces play a role in joint morphogenesis. However, how mechanical stimuli are translated into actual tissue growth and ultimately determine joint shape is still not well understood. Our computational model of joint morphogenesis provides a complementary tool to the experimental studies. Through hypotheses and simplifying assumptions, we have isolated critical contributors to the mechanotransduction of mechanical loading into chondrogenesis and subsequent shaping of the joint.

The computational results show that compressive fluid pressure is an adequate predictor of joint morphogenesis. In the predicted normalized surface growth map for the healthy growth case (Fig. 3E, top right) the ventral condyle exhibited a considerably larger amount of growth than the dorsal condyle, which is in agreement with the larger normalized area of the ventral condyles and no change of the dorsal condyles identified in the experimental control group with respect to the GSK101 group (Fig. 2F, top left). The predictions for the healthy growth case also exhibited more growth towards the distal area than the mechanically-impaired one (Fig. 3E, bottom right), which only had a slight gradient in the proximo-distal direction. These differences matched our experimental findings on decreased normalized volumes in both condyles of the GSK101 group with respect to the control group (Fig.2C, left).

Certainly, our model points to a relationship between the fluid pressure distribution and the shaping of the joint. Chondrocytes might not be sensing interstitial hydrostatic pressure directly, but rather a different biophysical factor related to it. Osmotic stresses have been repeatedly identified as the stimuli triggering a series of signalling events in relation to the TRPV4 channel, that are propagated into changes in gene expression and extracellular matrix synthesis. Yet, studies have shown that osmotic loading as well as mechanical loading elicit responses of the TRPV4 channel [23,27,28,30]. Recent publications suggest TRPV4 is a cell volume sensor and is activated regardless of the molecular mechanism underlying said volume change [67]. Furthermore, hydrostatic and osmotic pressures have similar effects on cartilage formation [68], and they both affect intracellular ion signalling in chondrocytes [69,70]. It is not within the scope of this study to determine the complex interrelations between the osmotic and hydrostatic pressures induced by mechanical loading on cartilage. Many studies have shown that hydrostatic pressure increases cartilage gene expression and extracellular matrix formation (see review by [71]) and, hence, we selected fluid pressure as a driver of mechanical growth in our model. Our computational results indicate that fluid pressure can predict local tissue growth in the experimentally-informed model of joint morphogenesis developed in this study.

### 4.4 Poroelasticity can be used to explore how dynamic loading dictates bone rudiment morphology

Due to the nature of the poroelastic tissue, only compressive dynamic loading can generate the non-homogeneous fluid pressure pattern within the humerus that dictates tissue growth in our computational model (Fig. 3B). In contrast, static loading generates an initial pressure distribution that quickly dissipates as fluid seeps out of the bone rudiment (SI Appendix, Fig. S9C). Such behaviour is in agreement with experimental studies showing that cartilage growth is promoted by repetitive compressive loading while static loading inhibits it [37–41]. Unlike our previous models of joint morphogenesis [34,35], we are now able to inherently capture the effect due to the type of loading imposed owing to the biphasic approach that incorporates the fluid flow component into the modelling. An earlier computational study [8] used poroelasticity to relate local patterns of biophysical stimuli to the emergence of joint shape in a model of a chick knee, but could not predict growth morphologies. Through the solid component growth, our model goes a step further and can more confidently relate local tissue growth to final bone rudiment morphology based on cyclic loading-induced mechanical stimuli.

We explored alternatives to the compressive pore pressure as mechanical stimuli for our growth model (SI Appendix, Figs. S10 and S11). Several measures of compression and fluid flow in the tissue were considered, with the idea of identifying and implementing an alternative mechanical growth stimulus in our formulation. We selected the positive divergence of the seepage velocity (SI Appendix, Fig. S11A, top row) because it is a measure of the rate of compression on the solid component of the material, and its distribution within the humerus is quite different from the fluid pressure pattern. The resulting local tissue growth due to the mechanical contribution was distributed more evenly towards the distal part of the humerus (SI Appendix, Fig. S13A), instead of being localized below the radius contact loading (Fig. 3C). In addition, less growth was observed in the proximal part of the humerus for the alternative model. Interestingly, this produced an apparent rotation of the humerus grown surface (SI Appendix, Fig. S13B) rather than the slight bending and outward growth observed broadly around the ventral condyle region for the pressure-based mechanical growth (Fig. 3D, top row). Further study would be required to ensure artefacts due to inadequate loading or boundary conditions are not at play here before discarding the rate of tissue compression as a potential biophysical stimuli within the joint morphogenesis process.

These exploratory simulations demonstrate the potential of the proposed model as a tool to unravel the mechanisms at play in the shaping of the joint. Through the computational study of how different measures of pressure, compression, and fluid flow evolve in response to loading setups representative of in vivo conditions, we could identify potential biophysical stimuli for further study in experiments.

### 4.5 Potential of combined experimental and computational approaches

Significant progress is being made in determining the activation mechanisms of the TRPV4 and Piezo mechanosensitive channels. Yet, the connection across scales – from organ-level loading to molecular response – is often overlooked. Through additional experimental studies that target mechanosensitive channels other than TRPV4, we could better distinguish the different mechanical stimuli involved in the shaping of the joint. Together with improved computational models, e.g. incorporating osmotic pressure into the formulation and modelling the complete joint, it would allow us to better identify the biophysical drivers of growth during joint formation.

We must expand our focus beyond the joint morphogenesis stage. Separation during cavitation has been suggested to rely on mechanically-induced changes in the extracellular matrix [72]. Mechanical forces are also known to affect the ossification process in the later stages of joint development [3]. Further studies are required to clarify the mechanosignalling mechanisms at play during these stages. Emerging experimental techniques like whole mount staining and imaging provide the opportunity to explore the 3D spatial distribution of mechanosensitive growth regulators [73] involved in cavitation and ossification. Developing such experiments in close association with computational modelling will provide a powerful tool to further explore the mechanobiology of joint formation.

## 5 Conclusions

The effect of loading-induced mechanical stimuli on joint morphogenesis was studied through the quantification of 3D humeri shapes in regenerating axolotl forelimbs. Normally-regenerating limbs were compared to those of animals in which chondrocyte mechanosensitivity was impaired by the administration of a TRPV4 agonist. Results demonstrated that mechanics has a role in the shaping of the joint.

We developed a finite element model of joint morphogenesis with cartilage modelled as a poroelastic material, in which growth of the solid part was due to a constant biological component as well as a loading-dependent mechanical component that includes its dynamic effects. Computational results indicated fluid pore pressure is a reasonable predictor of local tissue growth and ultimate joint shape, even if chondrocytes might not be directly sensing and responding to hydrostatic pressure. The computational model presented provides a tool to explore alternative mechanical stimuli that may also be critical in joint morphogenesis, such as static loading or constrained conditions.

Integrating experiments and computational modelling provides interesting insights that experiments alone cannot deliver. The combined approach presented in this work allowed us to validate the mechanical regulatory hypotheses with an in silico model. Such methodology will become indispensable as we advance in the study of mechanobiological processes like those involved in joint formation.

## Ethics

Animal experimentation: axolotls (*Ambystoma mexicanum*: d/d RRID Catalog #101L) were either bred in captivity at Northeastern University or purchased from the Ambystoma Genetic Stock Center at the University of Kentucky. Experiments were performed in accordance with Northeastern University Institutional Animal Care and Use Committee. For all experiments, animals were anaesthetised by treatment of 0.01% benzocaine until visually immobilized.

## Data Accessibility

The original microscopy data are available upon request. The scripts and some example files used in the experimental data analysis pipeline are available on the Zenodo repository, accessible at https://doi.org/10.1101/2021.08.28.458034. The computational code and simulation input files can be accessed at https://github.com/ecomellas/CompLimb-biomech.git. All other study data are included in the article and/or supporting information.

## Author Contributions

SJS and JRM conceived, designed and coordinated the study. JEF performed the main axolotl experiments and TJD provided additional experimental data. EC, GK, KL and TM processed the experimental data under the guidance of SJS. EC, SJS, JRM and TJD analysed and interpreted the experimental results. EC developed the theoretical model formulation with the collaboration of SJS and JJM. EC implemented the computational model and performed the numerical simulations. EC, SJS and JJM analysed and interpreted the computational results. EC and SJS drafted the manuscript. All authors revised and approved it for publication.

## Acknowledgments

The authors would like to acknowledge Eun Kyung Jeon for providing the far red nuclear stained limb image (SI Appendix), Vineel Kondiboyina for the nanoindentation test data (SI Appendix), Yash Kulkarni for contributing to the bone rudiment segmentations, and Soha Ben Tahar for exploratory finite element studies. EC thanks Markéta Tesařová for sharing their stl files of a newt limb.

Microscopy images were obtained from the Harvard University Center for Biological Imaging and the Northeastern University Chemical Imaging of Living Systems core. This work was completed using the Discovery cluster, supported by Northeastern University’s Research Computing team. We acknowledge animal support from the Ambystoma Genetic Stock Center funded by NIH grant P40-OD019794.

