## Supplementary Information for "Local mechanical stimuli shape tissue growth in vertebrate joint morphogenesis"

#### Contents

**S1 Suppl. figure: Experimental methods** **2**

**S2 Suppl. figure: Experimental data analysis** **3**

**S3 Suppl. figure: Individual 2D surface maps** **5**

**S4 Suppl. text: Numerical framework for poroelasticity with continuum growth** **6**

**S5 Suppl. figure: Finite element model of the humerus** **10**

**S6 Suppl. text: Parameters of the finite element model of joint morphogenesis** **11**

**S7 Suppl. figure: Computational predictions of potential mechanical stimuli** **15**

**S8 Suppl. text: Alternative measures of humeri shaft size** **17**

**S9 Suppl. figure: Alternative finite element growth model** **18**

**References** **19**

### S1 Suppl. figure: Experimental methods

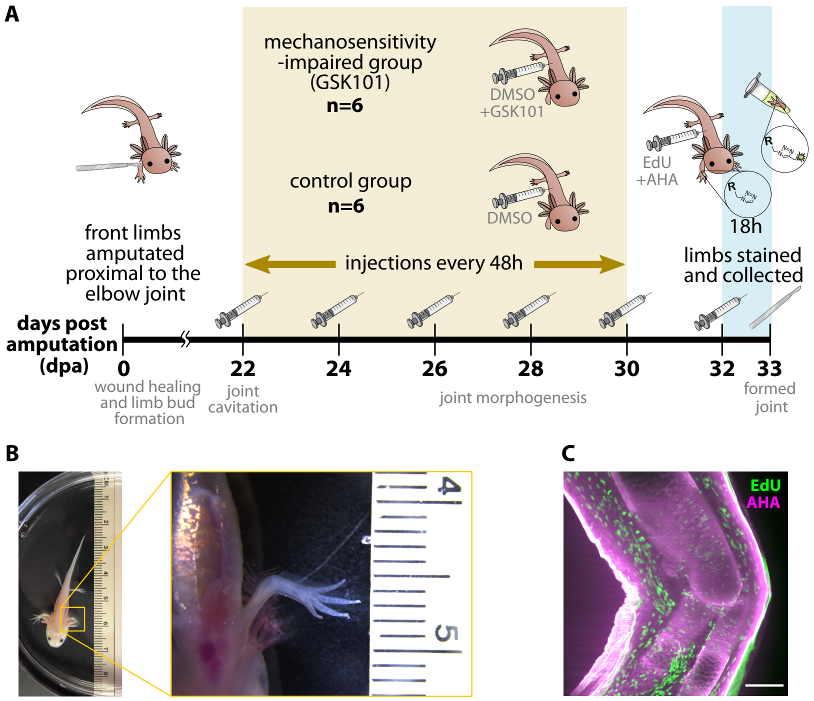

**Figure S1:** Overview of the experimental methods. (A) Illustrated timeline of the experiments. Created with images from `BioRender.com` and `smart.servier.com`. (B) Animals 3-5 cm in size were used, similar to the one shown here. Both the image of the whole animal as well as the close-up of a forelimb include a ruler in cm. (C) Axolotl forelimbs were imaged following the whole-mount click-it based visualization technique in [1] to obtain an image stack of the regenerated elbow joint. A central slice of a 3D image stack is shown here. The scale bar length represents 300 µm. Image acquired using a Zeiss light sheet Z.1 microscope paired with Zen software. In-plane pixel resolution is 0.9154 µm and slices are 4.9454 µm apart. The file size containing both the EdU and AHA channels is about 3 GB.

### S2 Suppl. figure: Experimental data analysis

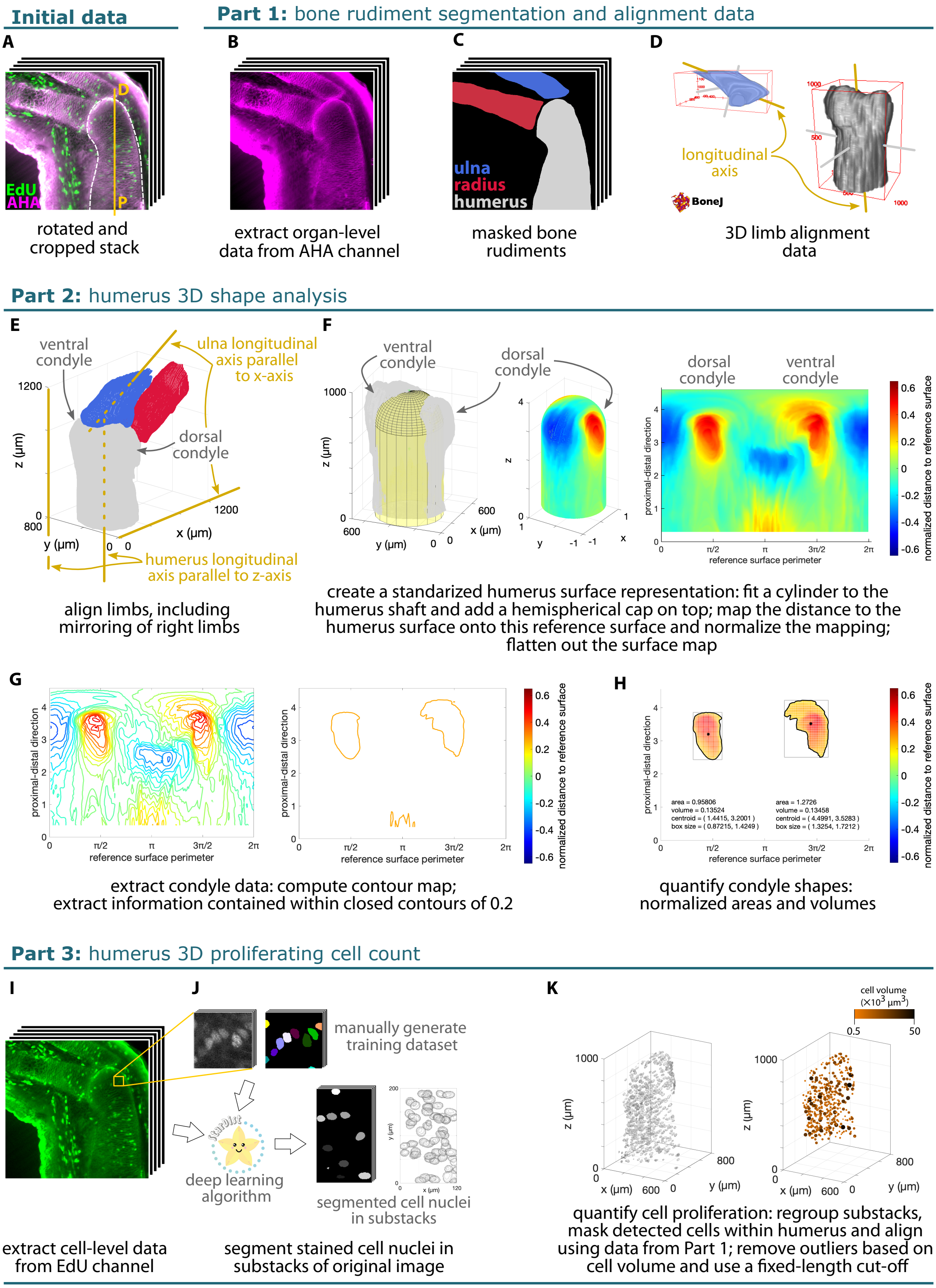

**Figure S2** ***(previous page)*****:** Complete workflow of the experimental data analysis using an exemplary control limb. (A) Each 3D image stack was cropped around the elbow joint and rotated to vertically align the proximo-distal (P-D) axis of the humerus. (B) The AHA staining in (A) allowed for segmentation of the bone rudiments, producing (C) the masks of the radius, ulna and humerus. (D) The Fiji plugin BoneJ [2, 3] provided data for limb alignment, based on the principal axes of the humerus and ulna bone rudiments. The minimum principal axis computed by BoneJ corresponded with the proximo-distal longitudinal axis of the bone rudiment. (E) Using data from (D), the surfaces in (C) were aligned in 3D space using Matlab [4]. (F) The aligned humerus from (E) was mapped onto a reference surface and normalized with the fitted cylinder diameter to create a 2D representation of the humerus' 3D surface. (G) The representation of the ventral and dorsal condyles were extracted and (H) quantified for each limb. (I) The EdU staining in (A) was used to identify the proliferating cell nuclei within the humerus bone rudiment. (J) Stardist3D [5] was trained with a custom-made dataset. Substacks of the original 3D image were fed to the algorithm, which provided the corresponding cell nuclei segmentations. (K) Substacks of cell nuclei segmentations were re-grouped, nuclei within the humerus were extracted using the corresponding mask from (C) and aligned in space using data from (D). After removal of outliers, the position of each nucleus' centre of mass and volume was plotted in 3D space. Total number of cells were counted within an equivalent volume among all humeri.

### S3 Suppl. figure: Individual 2D surface maps

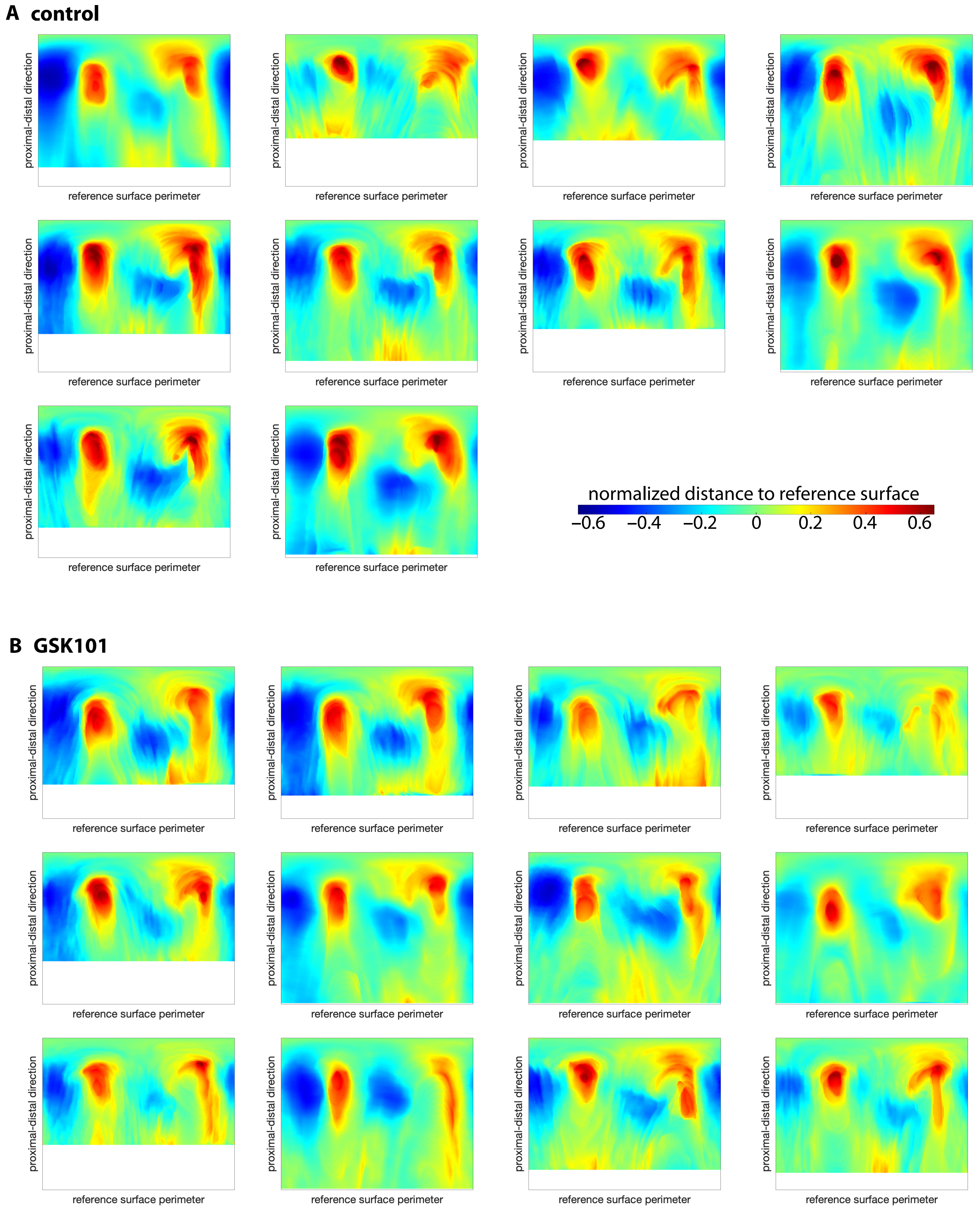

**Figure S3:** Individual 2D surface maps of the analyzed axolotl humeri for (A) the control group and (B) the GSK101-treated animals.The 2D surface maps of all humeri were computed following the methodology described in the main text. Maps have varying heights due to length variability in the segmented humerus bone rudiments. All maps have been aligned in the horizontal direction based on the centroid position of the dorsal condyle. Two control limbs had to be discarded because the segmented humerus was too short to be properly aligned following the methodology developed.

### S4 Suppl. text: Numerical framework for poroelasticity with continuum growth

We propose a finite element biomechanical model of growth at tissue level to study how specific changes in limb motion regulate joint morphology. Bone rudiments undergoing the joint morphogenesis stage are mostly composed of chondrocytes. Hence, the cartilaginous tissue is modeled as a biphasic poroelastic material consisting in a fluid-saturated nonlinear porous solid. The existing nonlinear poroelastic formulation [6] implemented in `deal.II` [7] was extended to incorporate continuum growth [8]. Here, we provide a brief overview of the poroelastic formulation and describe the derivation and implementation of the continuum growth portion of the computational model.

#### S4.1 Kinematics

The biphasic material is composed of a hyperelastic solid skeleton (S) and a pore fluid constituent (F) that occupy simultaneously a given spatial position $\boldsymbol{x}$ in the current configuration at time $t$. Then, the constituent deformation map is $\boldsymbol{x} = \chi_S(\boldsymbol{X}_S, t) = \chi_F(\boldsymbol{X}_F, t)$, where $\boldsymbol{X}_S$ and $\boldsymbol{X}_F$ correspond to the different material positions in the reference configuration at the reference time $t_0$ of the solid and fluid constituents, respectively. The solid displacement is $\boldsymbol{u}_S = \boldsymbol{x} - \boldsymbol{X}_S$, and its material deformation gradient is $\boldsymbol{F}_S = \partial\boldsymbol{x}/\partial\boldsymbol{X}_S$, where the subscript '$S$' will be dropped for clarity in the subsequent derivations. The seepage velocity describes the motion of the fluid with respect to the deforming solid material, i.e. $\boldsymbol{w}_F = \boldsymbol{v}_F - \boldsymbol{v}_S = \partial\chi_S/\partial t - \partial\chi_F/\partial t$.

Note that the solid and fluid constituents are assumed to be separately incompressible, but the biphasic material is compressible owing to the fluid flow within the pores of the deforming solid skeleton. In addition, the saturation condition establishes $n^S + n^F = 1$, where $n^S$ and $n^F$ are the volume fractions of the solid and fluid constituents, respectively. Based on the volume balance of the solid skeleton, the former can be integrated towards $n^S = n^S_{0S}/J$, where $J = \det(\boldsymbol{F}) > 0$ and $n^S_{0S}$ is the initial solid volume fraction, a measure of the biphasic material's initial porosity.

#### S4.2 Continuum growth

We introduce volumetric tissue growth through the multiplicative decomposition of the material deformation gradient tensor of the solid component (Fig. S4),

$$\boldsymbol{F} = \boldsymbol{F}^e \cdot \boldsymbol{F}^g \quad \text{with} \quad \boldsymbol{F}^g = \vartheta\boldsymbol{1}. \tag{1}$$

The rate of the growth stretch $\vartheta$ determines the isotropic growth, and $J^e = \det(\boldsymbol{F}^e) > 0$ is the Jacobian determinant of the elastic part of the material deformation gradient tensor of the solid component, $\boldsymbol{F}^e$.

Following past studies [9, 10], we hypothesize that growth stretch rate is given by the sum of a biological and a mechanical contributions,

$$\dot{\vartheta} = \dot{\vartheta}_b + \dot{\vartheta}_m. \tag{2}$$

The rationale is that certain growth will occur during limb formation regardless of external mechanical stimuli, owing to morphogenetic cues that result, for example, in cell proliferation and extracellular matrix deposition. The amount of biological growth is, therefore, assumed to be proportional to chondrocyte density, $C_d$. We experimentally measured chondrocyte density in a regenerated axolotl humerus and found that it was approximately constant along its proximo-distal axis (Fig. S5). For this reason, we used a constant biological growth function,

$$\dot{\vartheta}_b\,(C_d) = k_b, \tag{3}$$

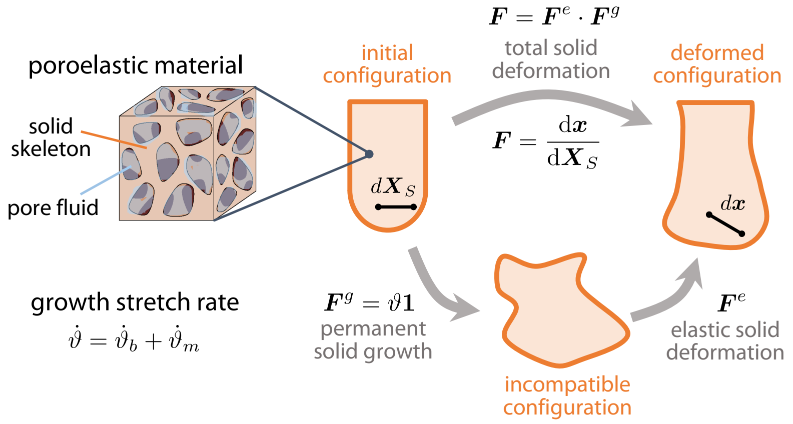

**Figure S4:** Continuum growth in the computational model is based on the multiplicative decomposition of the deformation gradient tensor $\boldsymbol{F}$ that characterizes the solid component of the poroelastic material. $\boldsymbol{F}$ maps a vector from the initial or reference configuration $d\boldsymbol{X}_S$ into a new position after deformation in the current configuration $d\boldsymbol{x}$. It is split into an elastic deformation gradient tensor $\boldsymbol{F}^e$ and a growth tensor $\boldsymbol{F}^g$. For simplicity, growth is assumed to be volumetric and proportional to the growth stretch variable $\vartheta$, whose rate is the sum of a biological contribution $\dot{\vartheta}_b$ and mechanical contribution $\dot{\vartheta}_m$.

where the parameter $k_b$ modulates the rate of tissue growth due to intrinsic biological factors. The mechanical contribution is proportional to the biophysical stimuli $\Xi$, hypothesized to drive local tissue growth. For simplicity, we started our numerical explorations assuming the positive (compression) pore pressure was driving the mechanical portion of tissue growth,

$$\dot{\vartheta}_m(\Xi) = k_m\,\Xi = k_m < p >, \tag{4}$$

where the parameter $k_m$ adjusts the proportion of mechanical growth to the overall tissue growth, and the Macaulay brackets $< \bullet >$ indicate that only positive (compressive) fluid pressure produces mechanical growth. We note that in addition to $\Xi \; =< \; p \; >$, and in order to explore different feedback mechanisms, other alternative mechanical stimuli have been tested, such as $\Xi =< \mathrm{div}\,(\boldsymbol{w}) >$, which is a measure of local solid component compression rate.

The algorithm used to implement continuum growth in the poroelastic model is given in Fig. S6.

#### S4.3 Governing field equations

Following standard assumptions as described in [6], the governing field equations are derived from the mass continuity and local linear momentum balance equations of the individual material components. The resulting weak form of the overall linear momentum balance is

$$\int_{\mathcal{B}_0} \nabla(\delta\boldsymbol{u}) : \boldsymbol{\tau}\,\mathrm{d}V_{0S} - \int_{\partial\mathcal{B}_0^T} \delta\boldsymbol{u}\cdot\boldsymbol{T}^*\,\mathrm{d}A_{0S} = 0 \qquad \forall\delta\boldsymbol{u}, \tag{5}$$

and the overall mass balance is

$$\int_{\mathcal{B}_0} \delta p\,\dot{J}_e\,\mathrm{d}V_{0S} - \int_{\mathcal{B}_0} \nabla(\delta p)\cdot\boldsymbol{w}\,J\,\mathrm{d}V_{0S} = 0. \qquad \forall\delta p. \tag{6}$$

Both equations are given in the reference configuration and introduce the solid displacement test function $\delta\boldsymbol{u}$ and the fluid pore pressure test function $\delta p$, respectively. The Kirchhoff stress tensor $\boldsymbol{\tau}$ in (5) is defined by the constitutive equation of the solid component (see next section) and $\boldsymbol{T}^*$ is the prescribed traction on the boundary $\mathcal{B}_0^T$. Here, we have neglected volumetric forces due

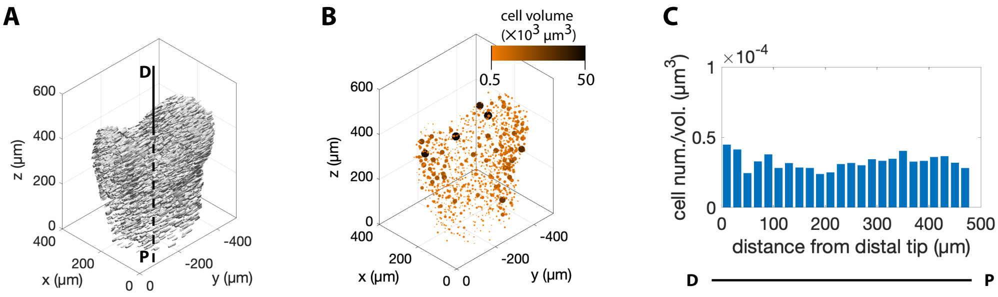

**Figure S5:** Experimental data used to determine a constant biological growth function. (A) The chondrocyte nuclei outlines were extracted from a far red nuclear staining 3D light sheet image stack of a regenerated axolotl forelimb using Cellpose [11]. Cell nuclei surfaces are shown in 3D space after vertically aligning the proximo-distal (P-D) axis of the humerus in Matlab [4]. (B) Cell nuclei positions and corresponding volumes in 3D space were obtained with the Fiji plugin 3D Objects Counter [12], imported and aligned in Matlab. Outliers were removed based on the cut-off volumes $0.5 \times 10^3$ and $50 \times 10^3$ µm. (C) Cell density was computed as cell number divided by cross-sectional slice volume, and plotted along the proximo-distal axis. A thickness of 20 µm was used to compute the humeri cross-sectional slice volumes from a segmentation of the whole humerus bone rudiment based on the original 3D image stack.

to the effect of gravity. The volume-weighted seepage velocity $\boldsymbol{w} = n^F \boldsymbol{w}_F$ introduced in (6) is defined by the constitutive equation of the fluid. The term $\dot{J}_e$ indicates the material time derivative of the Jacobian determinant of the elastic deformation gradient tensor. Here, we do not prescribe forced fluid flow across the boundaries.

#### S4.4 Constitutive models

The hyperelastic solid behavior is given by the constitutive equation

$$\boldsymbol{\tau} = \boldsymbol{\tau}_E^{\mathrm{NH}} + \boldsymbol{\tau}_E^{\mathrm{vol}} - p\, J\, \mathbf{1}, \tag{7}$$

where the ‘extra’ stress is split into a neo-Hookean contribution,

$$\boldsymbol{\tau}_E^{\mathrm{NH}} = \mu \left[ \boldsymbol{F}_e \cdot \boldsymbol{F}_e^T - \mathbf{1} \right], \tag{8}$$

and a volumetric term, which accounts for the compressibility effects of the biphasic material,

$$\boldsymbol{\tau}_E^{\mathrm{vol}} = \lambda \left[ 1 - n_{0S}^S \right]^2 \left[ \frac{J_e}{1 - n_{0S}^S} - \frac{J_e}{J_e - n_{0S}^S} \right] \mathbf{1}. \tag{9}$$

Here, we introduce the neo-Hookean shear modulus $\mu$ and the first Lamé parameter $\lambda$.

A Darcy-like law is used to define the fluid constitutive behavior,

$$\boldsymbol{w} = -\frac{1}{\mu^{\mathrm{FR}}} \left[ \frac{J - n_{0S}^S}{1 - n_{0S}^S} \right] \boldsymbol{K}_0^S \cdot \nabla p, \tag{10}$$

where, for simplicity, gravity contributions have been neglected. Here, we introduce the effective shear viscosity of the fluid, $\mu^{\mathrm{FR}}$ and the initial intrinsic permeability $\boldsymbol{K}_0^S = K_0 \mathbf{1}$, which is assumed to be isotropic.

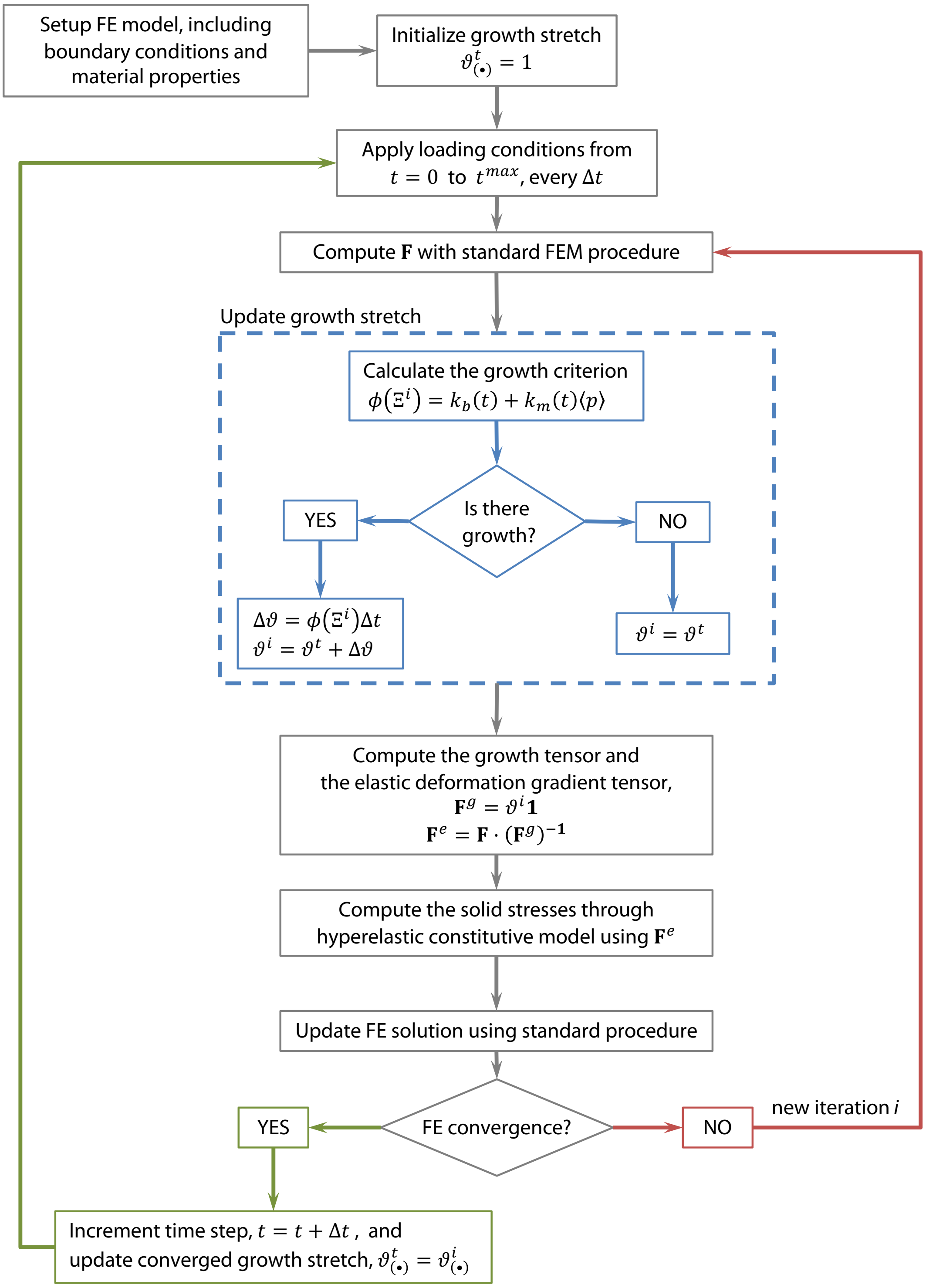

**Figure S6:** Algorithm for the numerical implementation of growth in the poroelastic finite element formulation used. Adapted from [8], the growth stretch in this study does not have a limiting function. The growth increment $\Delta\vartheta$ can be computed directly because the growth criterion is independent of previous values of $\vartheta^t$. The growth rate parameters $k_b$ and $k_m$ are time step dependent to ensure no growth is applied in the first and last time steps, corresponding to the application and removal of the loading conditions.

### S5 Suppl. figure: Finite element model of the humerus

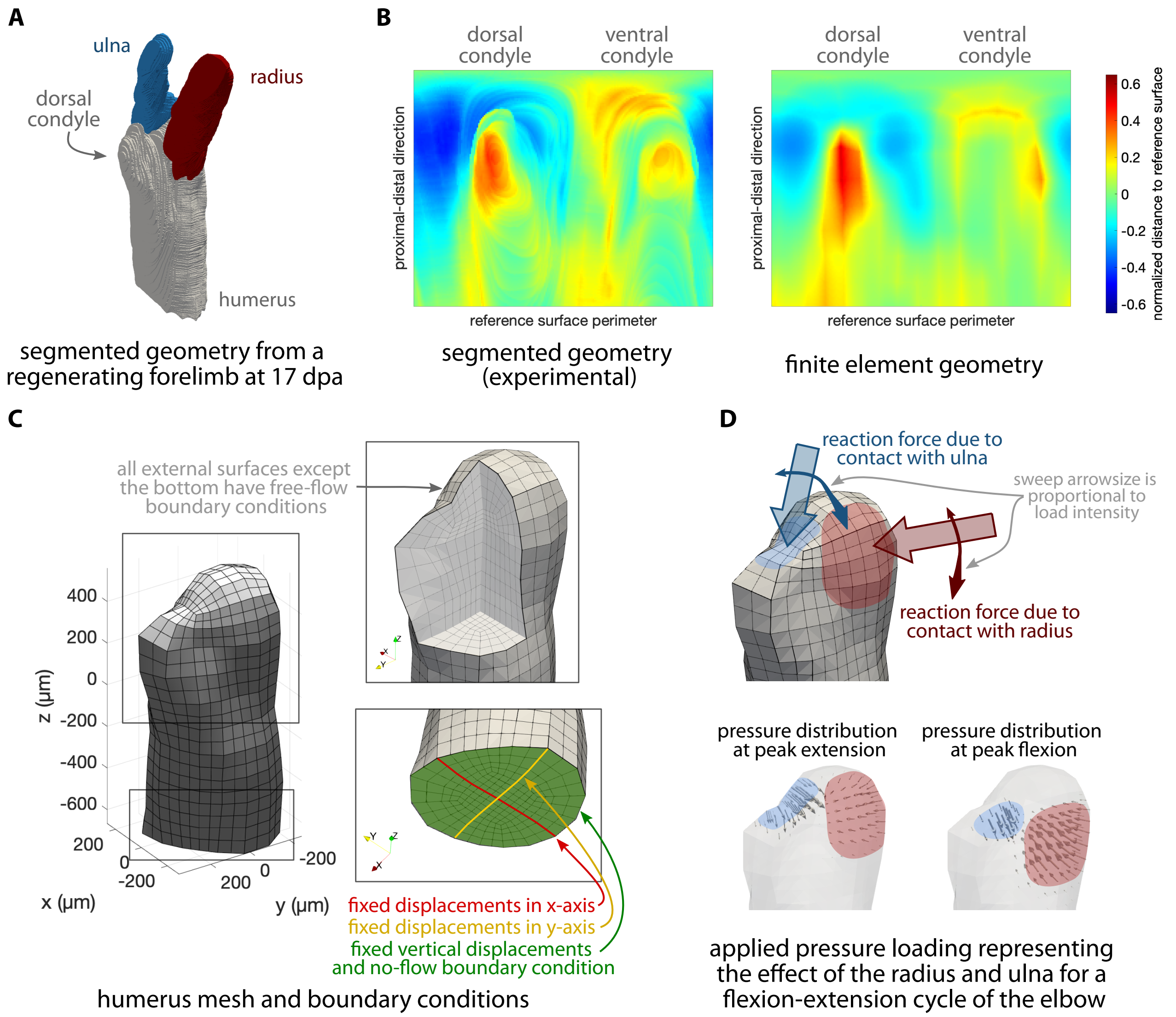

**Figure S7:** Finite element model of the humerus. (A) Segmentation of a regenerating forelimb at 17 days post-amputation (dpa) used as basis for the geometry. (B) 2D surface maps of the segmented humerus geometry and corresponding finite element approximation are similar, confirming the computational geometry is a good approximation to the experimental humerus surface. (C) Meshed humerus and boundary conditions applied in the computational simulations. (D) Loading to simulate a flexion-extension cycle of the elbow was applied as a sweeping motion together with a twofold increase in pressure load intensity at peak flexion.

### S6 Suppl. text: Parameters of the finite element model of joint morphogenesis

The material parameters used in the computational simulations are summarized in Table S1. In the definition of the solid component behavior, we selected the neo-Hookean shear modulus and the initial solid volume fraction based on values found in literature [13, 14]. The first Lamé parameter was set to a value roughly two orders of magnitude higher than the shear modulus to ensure correct enforcement of the compaction point behavior and of the incompressibility of the solid component. Given the small loading values applied in our simulations, predicted deformations throughout our model were always far from this point and, hence, this value does not impact the predicted patterns. The fluid component behavior was defined through the initial intrinsic permeability and the fluid viscosity. Preliminary simulations revealed that the value of the former had a considerable impact on the predicted pressure patterns. Therefore, initial intrinsic permeability was estimated based on experimental data, as explained below. The fluid viscosity was set to that of water at 25°C. Finally, the parameters regulating the contribution of the mechanical and biological growth rates to the overall tissue growth were manually adjusted to obtain a reasonable proportion between the two contributions for the healthy growth case.

A loading pressure range between 10 kPa at peak extension and 20 kPa at peak flexion was applied. This value is a rough guess based on the maximum muscle stress reported for tiger salamanders [15] (obtained for an animal mass of 1.3 g, which is the mean mass of the axolotls in our study) and an extrapolation of the relative cross-sections between the limb muscle and bone rudiment morphologies of a Spanish ribbed newt [16]. The increase in load intensity can be interpreted as a representation of the changes in contact force between the bone rudiments due to muscle contractions driving the limb flexion-extension motion. Contact area and sweep path were estimated based on the bone rudiment 3D surfaces extracted from experimental data.

Initially, the whole external surface of the humerus was set to allow free fluid flow across it. Upon close analysis of preliminary results, we considered that a bottom surface with no-flow condition approximated better the in vivo conditions of the humerus bone rudiment (Fig. S8). We simulated the ulna and radius contact separately with the goal of discerning the contribution of each load to the predicted patterns. For each simulation we considered only the sweeping motion without any load intensity change and, then, a fixed position but a load intensity change. Growth predictions for a free-flow and a no-flow bottom surface are roughly equivalent in the distal portion of the humerus for any given loading condition. Close to the bottom surface, differences in growth appear due to different pressure patterns predicted for the free-flow vs no-flow conditions. A free-

**Table S1:** Material parameters used in the computational simulations.

| Parameter | Symbol | Value | Units |
|---|---|---|---|
| solid shear modulus | $\mu$ | 2.0e3 | kPa |
| first Lamé parameter | $\lambda$ | 1.0e5 | kPa |
| initial solid volume fraction | $n_{0S}^{S}$ | 0.17 | |
| initial intrinsic permeability | $K_0$ | 1.0e-3 | $\mu$m$^2$ |
| fluid viscosity | $\mu_{\mathrm{FR}}$ | 0.89 | kPa·s |
| biological growth rate parameter | $k_b$ | 2.0e-5 | s$^{-1}$ |
| mechanical growth rate parameter | $k_m$ | 1.0e-5 | (kPa·s)$^{-1}$ |

flow bottom boundary condition enforces all nodes in the surface to have a zero pressure value, which prevents pressure build up above the surface. Ideally, we would want to avoid artefacts like these, which arise from a fictitious boundary, by modeling the complete bone rudiment. However, due to computational limitations and because we are interested in the growth of the distal portion of the humerus, we deemed that the approximation provided by our model was sufficient for the purposes of this study.

This set of simulations also served to confirm that the contribution of the radius loading to growth is much larger than that of the ulna loading. In our main simulations we combined sweeping motion with load intensity increment for both ulna and radius loading. In this way we simulated the change in contact position between bone rudiments during a flexion-extension movement (sweeping motion) as well as changes in reaction force between bone rudiments in contact owing to the effect of muscles contracting (intensity change).

Time steps of 0.01s were applied for a total of 1.01s to reproduce a flexion-extension cycle. Load value increased in a sinusoidal manner, as did the sweeping motion where the load was applied. The use of sinusoidal increments avoids abrupt changes in the numerical simulation, which are known to produce unrealistic peaks in predicted variable values. In fact, this is why growth was not applied in the first and last step, when the load was applied and removed. Preliminary simulations were run to ensure time step size was not affecting the predicted outcomes. We observed comparable pressure patterns for simulations with same material parameters, boundary conditions and loading patterns.

We used a stress-relaxation model to identify the initial intrinsic permeability value, $K_0 = 10^{-2}$ µm in which computational predictions were compared to experimental data from a nanoindentation test on an axolotl bone rudiment. The indentation rate was 100 nm/s and the maximum indentation distance of 5 µm was applied with an indenter of radius 25 µm. Relaxation was recorded over a 3-minute period. We predicted a similar stress-relaxation condition in our material by applying a static load on the distal part of the humerus (Fig. S9A). We then measured the total reaction force of the loaded surface over time for different values of $K_{0s}$ and compared them to the experimental results (Fig. S9B). This computational example illustrates the advantage of using a poroelastic material model instead of an elastic one. We are able to explicitly capture the relaxation of the material under a static load. In this way, we can predict (and adjust) the dissipation of the pressure accumulated below the loading surface as fluid seeps out of the sample and the material returns to its relaxed state (Fig. S9C).

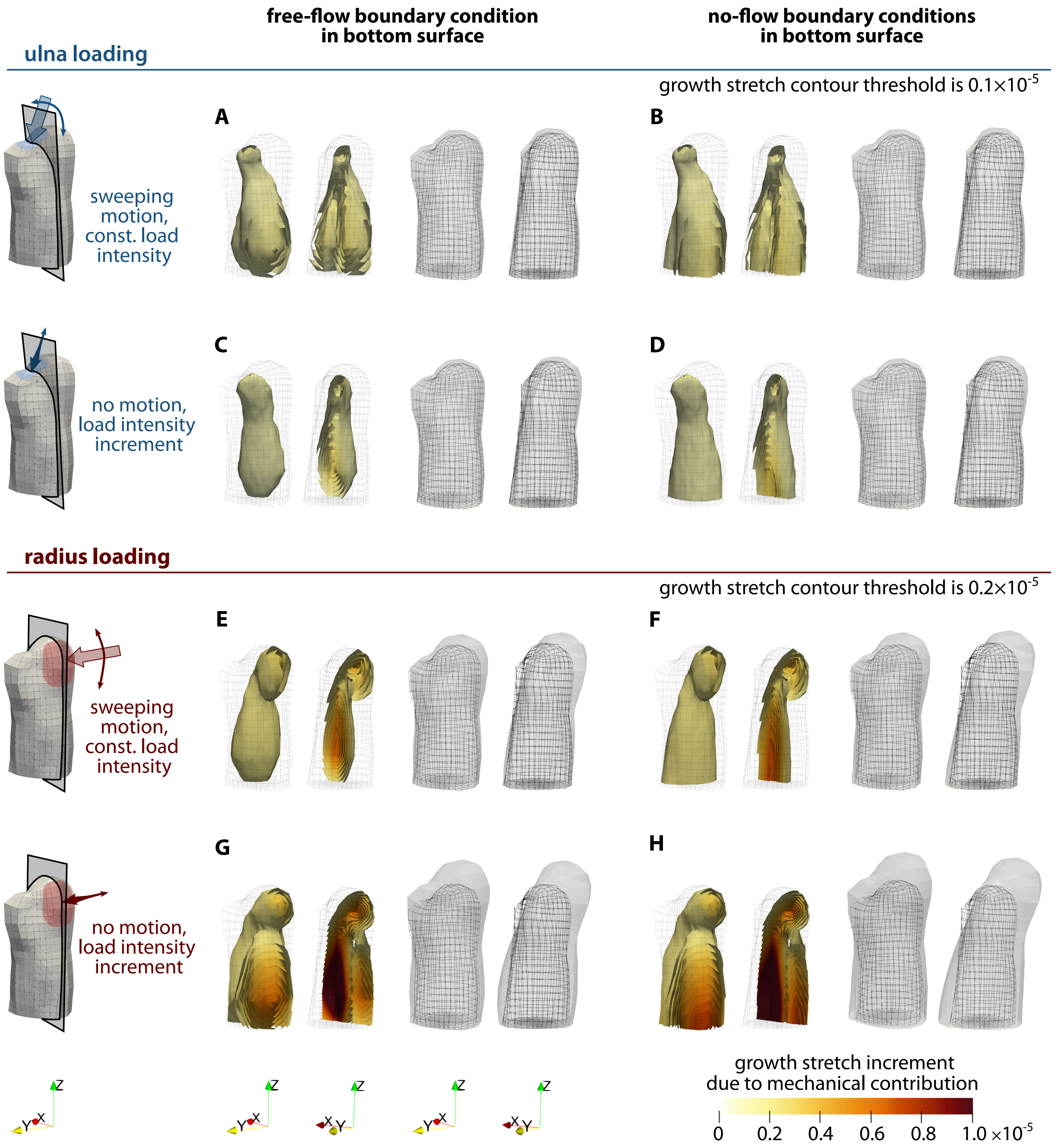

**Figure S8:** Computational predictions for free-flow and no-flow boundary conditions in the bottom surface. Results are shown separately for ulna (A-D) and radius (E-H) loading, with sweeping motion (A,B,E,F) and load intensity change (C,D,G,H) also studied separately. For each loading and boundary conditions combination the left-most image shows the growth stretch increment distribution in the humerus and the centre left image shows a clipped view of the same result. Growth shown is exclusively due to the mechanical contribution, with $k_m = 5 \cdot 10^{-4}$ (kPa·s)$^{-1}$ at the end of a flexion-extension load cycle. The centre right and right-most images of each simulation correspond to the frontal and side views of the grown humerus shape. For all cases, growth computed after a 1-second-cycle is scaled by a factor of 36000, representing 10 hours of loading.

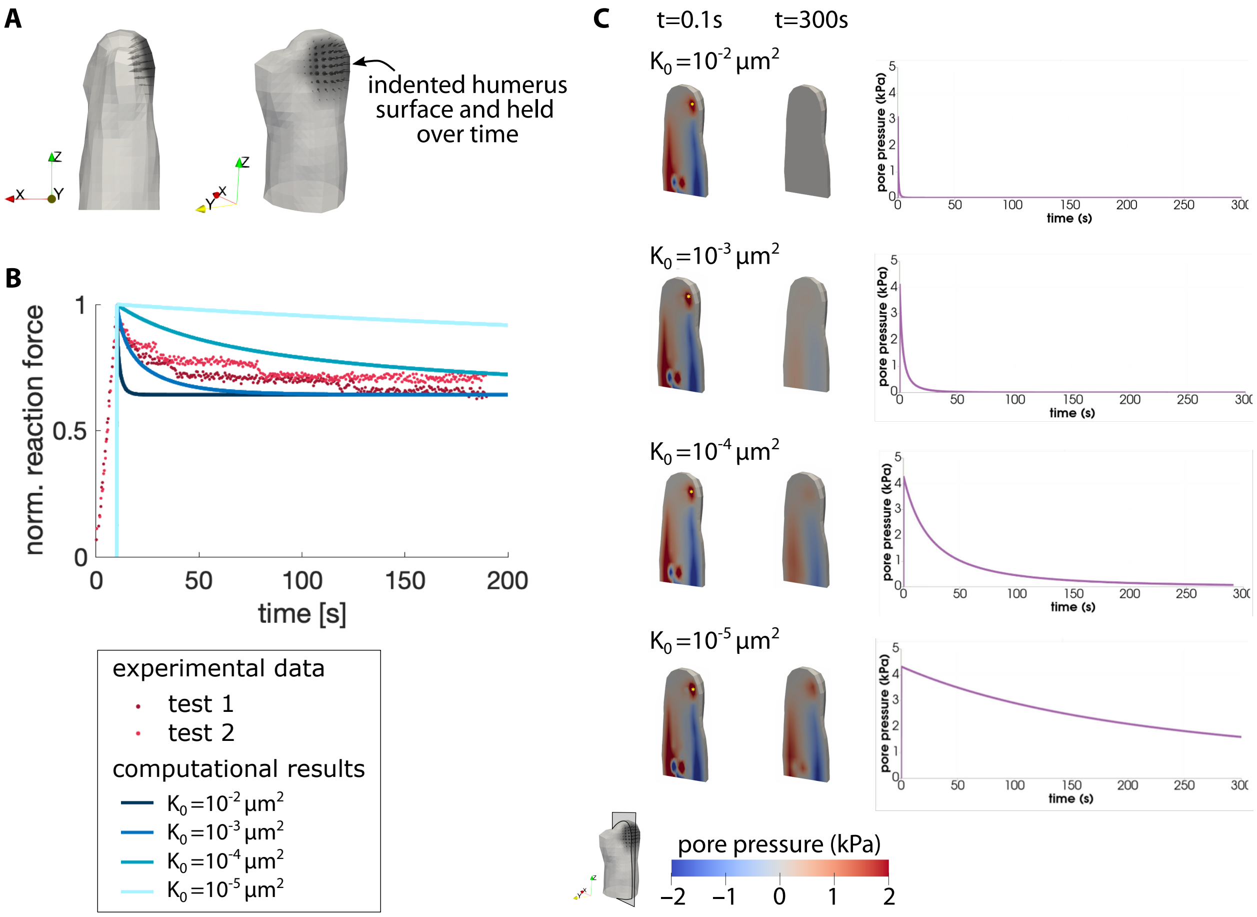

**Figure S9:** Adjustment of the initial intrinsic permeability based on experimental data and using non-cyclic constant indentation. (A) Stress-relaxation computational model used. (B) Reaction force obtained in the nanoindentation experiments on an axolotl limb bone rudiment (dots) and reaction force computed in the stress-relaxation model for different values of $K_0$ (lines). Reaction force is normalized with its peak value over time for each example. (C) Pressure distribution at the initial (left column) and final (middle column) time steps in a a vertical cross-section of the humerus (shown bottom left) for the different values of $K_0$. The evolution of the pressure value at the point marked with a yellow asterisk in the left column is shown over time in the right column.

163 # S7 Suppl. figure: Computational predictions of potential
164 mechanical stimuli

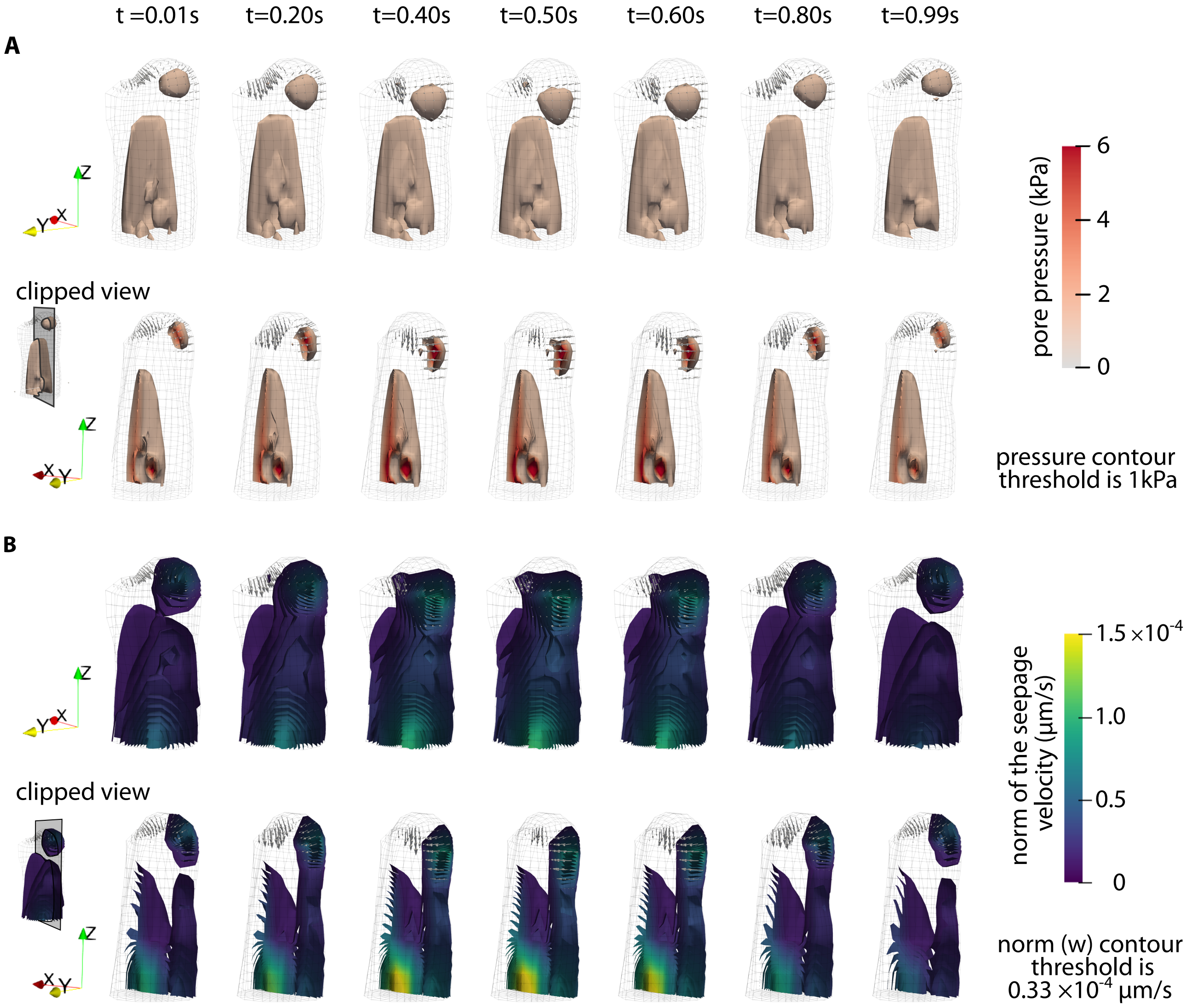

**Figure S10:** Computational predictions of potential mechanical stimuli that chondrocytes sense and respond to, and could be driving joint morphogenesis. The evolution of patterns within the humerus bone rudiment is shown over a 1-second flexion-extension cycle. (A) Predicted pore pressure distribution in the whole humerus (top row) and in a clipped view (bottom row), both given in kPa. (B) Distribution of the L2-norm of the seepage velocity in the humerus (top row) and in a clipped view (bottom row), both given in mm/s. The volume-weighted seepage velocity $\boldsymbol{w}$ is the relative velocity of the pore fluid with respect to the deforming solid skeleton. Its L2-norm is a measure of the amount of fluid flow. The general pattern of the norm of the seepage velocity (B) resembles the predicted pressure patterns (A) and would probably result in similar growth predictions.

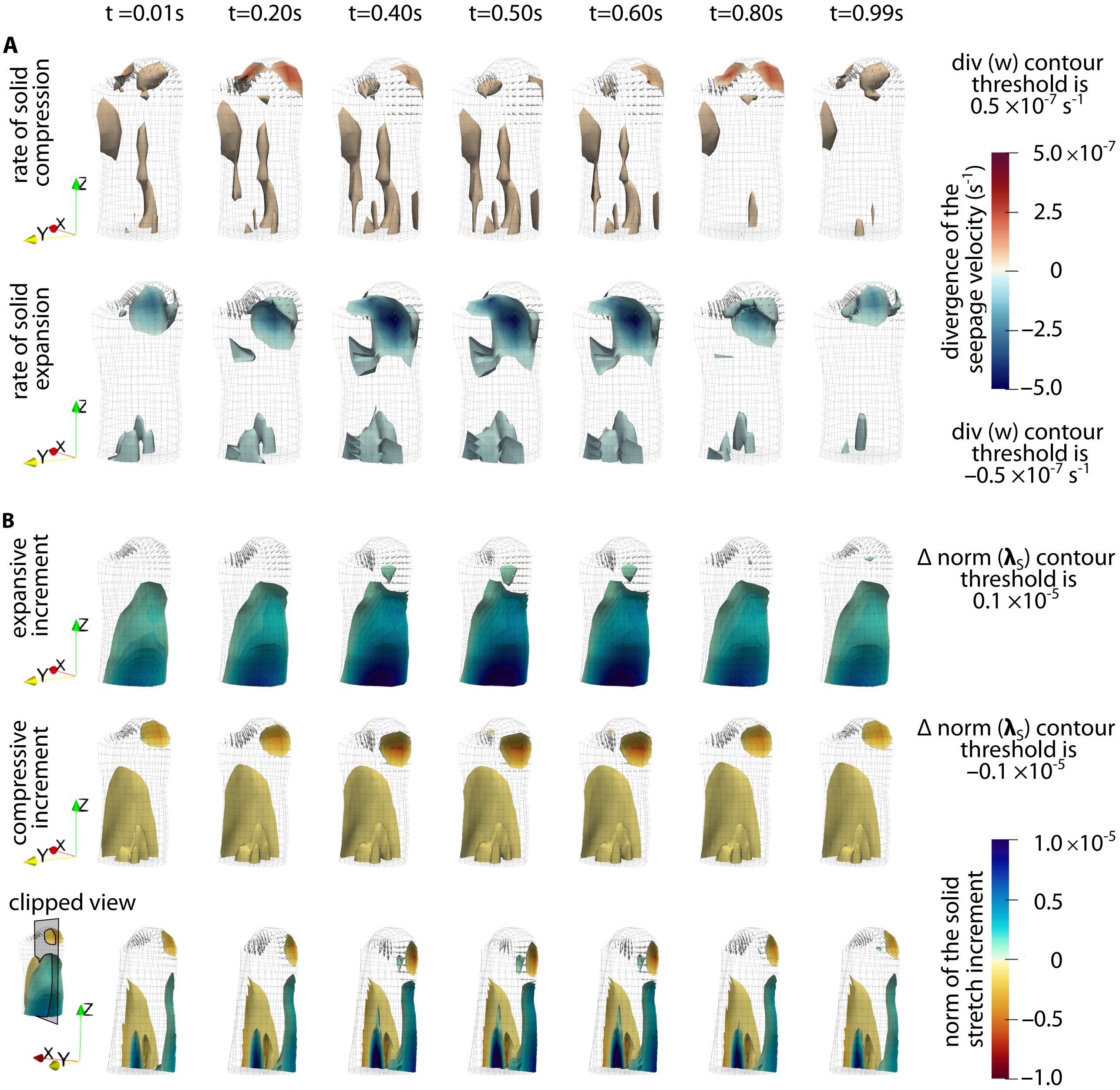

**Figure S11:** Computational predictions of potential mechanical stimuli that chondrocytes sense and respond to, and could be driving joint morphogenesis (continued). The evolution of patterns within the humerus bone rudiment is shown over a 1-second flexion-extension cycle. (A) Distribution of the divergence of the seepage velocity, given in $s^{-1}$. A positive value indicates rate of solid compression (top row) while a negative one indicates rate of solid expansion (bottom row). (B) Distribution of the increment of a scalar norm of the solid stretch, computed as $\Delta\mathrm{norm}(\boldsymbol{\lambda}_S) = \sqrt{(\lambda_x^2 + \lambda_y^2 + \lambda_z^2)/3} - 1$, where $\lambda_x$, $\lambda_y$ and $\lambda_z$ are the stretches of the solid in the global coordinates. A positive increment of $\Delta\mathrm{norm}(\boldsymbol{\lambda}_S)$ indicates solid expansion (top row) while a negative one indicates solid compression (middle row). The two are superimposed in a clipped view of the humerus (bottom row). The compressive increment of the norm of the solid stretches (B, center row) resembles the predicted pressure patterns (Fig. S10A) and would probably result in similar growth predictions. However, the rate of compression (A, top row) produces a noticeably different pattern. Fig. S13 shows the predicted growth for this mechanical stimuli.

### S8 Suppl. text: Alternative measures of humeri shaft size

Our statistical analysis of the diameters of the cylinders fitted to the humeri shaft using the Matlab [4] File Exchange function ‘cylinderfit’ produced no significant difference between the control and GSK101-treated groups (Main Text, Fig. 2B). To ensure this result was not due to an insufficiently sensitive method of measurement of the humeri shaft size, we computed additional metrics using an alternative methodology.

First, we extracted a standardized portion of the humerus shaft by trimming the distal and proximal parts of the aligned humerus surface (gray surface in Main Text, Fig. 1D). We removed the surface portion above a vertical distance equal to the fitted diameter (measured from the most distal part along the vertical axis), and then kept a portion of the shaft equal to 0.25 times the fitted diameter in thickness. To measure the size of the extracted shaft, it was divided into 20-µm-thick slices and the surface of each slice was projected onto the cross-sectional x-y plane. The resulting 2D projected shape was converted to a binary image. Given that the extracted shafts were about 120 µm in thickness, we obtained between 5 and 7 projected 2D shapes per humerus. Finally, we computed a series of shape metrics for each projected shape and averaged the values of each for all shapes in a limb. The metrics computed were area, perimeter, equivalent diameter (i.e. the diameter of a circle with the same area as the projected shape), major axis, and minor axis. Results confirm that there is no significant difference in humeri size between the control and GSK101-treated groups (Fig. S12).

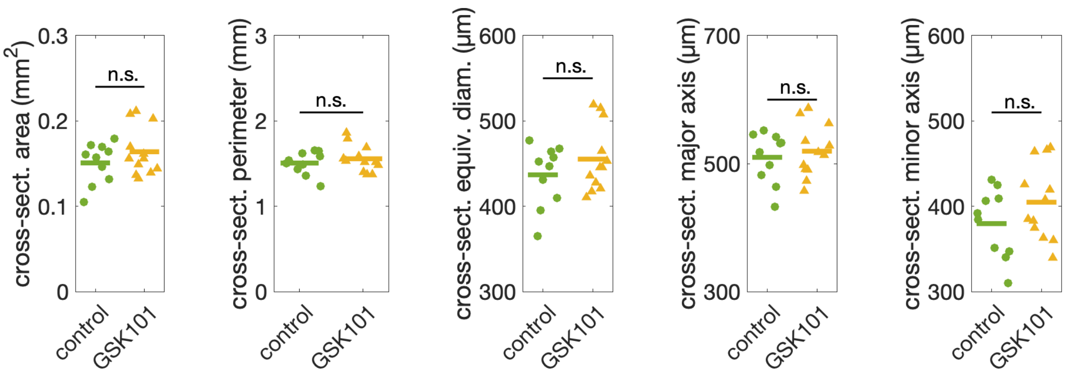

**Figure S12:** Results of the statistical analysis on the data points obtained following an alternative methodology to measure humeri shaft size. All data was normally distributed (Shapiro-Wilk test) and we performed a one-way ANOVA test to obtain p-values, which were all above 0.1.

### S9 Suppl. figure: Alternative finite element growth model

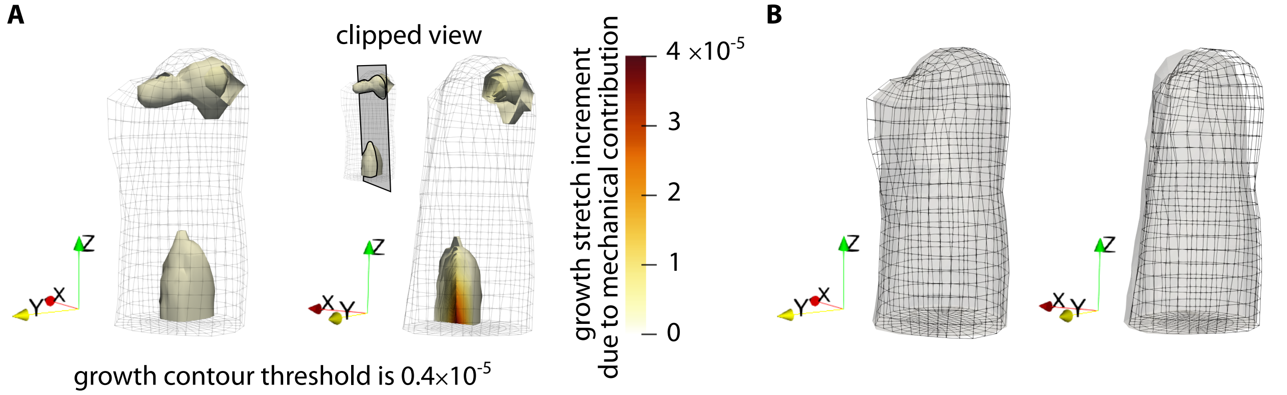

**Figure S13:** Finite element growth model using $< \mathrm{div}\,(\boldsymbol{w}) >$ (positive divergence of the seepage velocity, a measure of the rate of solid compression) as mechanical stimulus. (A) Computational predictions of the local tissue growth due to the mechanical contribution at the end of one flexion-extension cycle. Distribution in the whole humerus (left) and in a clipped view (right). (B) Grown humerus shape scaled by a factor of 86400, representing 24 hours of loading. A frontal view (left) and a side view (right) are shown. A value of $k_m = 10$ was used to obtain local tissue growth of the same order of magnitude as in our previous simulations. The grown humerus shape showed considerably less amount of surface growth and appeared to rotate instead of bend like in the pressure-driven mechanical growth predictions. Further studies would have to be conducted to explore the effect of loading and boundary conditions on this model.

[4] MATLAB. version 9.7.0.813654 (R2019b). Natick, Massachusetts: The MathWorks Inc.; 2019.
